# How thalamic relays might orchestrate supervised deep training and symbolic computation in the brain

**DOI:** 10.1101/304980

**Authors:** Kenneth J. Hayworth, Adam H. Marblestone

## Abstract

The thalamus appears to be involved in the flexible routing of information among cortical areas, yet the computational implications of such routing are only beginning to be explored. Here we create a connectionist model of how selectively gated cortico-thalamo-cortical relays could underpin both symbolic and sub-symbolic computations. We first show how gateable relays can be used to create a Dynamically Partitionable Auto-Associative Network (DPAAN) (Hayworth, 2012) consisting of a set of cross-connected cortical memory buffers. All buffers and relays in a DPAAN are trained simultaneously to have a common set of stable attractor states that become the symbol vocabulary of the DPAAN. We show via simulations that such a DPAAN can support operations necessary for syntactic rule-based computation, namely buffer-to-buffer copying and equality detection. We then provide each DPAAN module with a multilayer input network trained to map sensory inputs to the DPAAN’s symbol vocabulary, and demonstrate how gateable thalamic relays can provide recall and clamping operations to train this input network by Contrastive Hebbian Learning (CHL) (Xie and Seung, 2003). We suggest that many such DPAAN modules may exist at the highest levels of the brain’s sensory hierarchies and show how a joint snapshot of the contents of multiple DPAAN modules can be stored as a declarative memory in a simple model of the hippocampus. We speculate that such an architecture might first have been ‘discovered’ by evolution as a means to bootstrap learning of more meaningful cortical representations feeding the striatum, eventually leading to a system that could support symbolic computation. Our model serves as a bridging hypothesis for linking controllable thalamo-cortical information routing with computations that could underlie aspects of both learning and symbolic reasoning in the brain.

## Introduction

In this paper, we propose a theory for how the biological brain might integrate trainable computation, analogous to learning in deep neural networks (Marblestone et al., 2016; Roelfsema and Holtmaat, 2018), with classical symbolic computation, analogous to “production system” models of cognition (Newell, 1994; Taatgen and Anderson, 2010), inside a single network architecture. We suggest that relay of information from a source memory buffer in the cortex, through the thalamus, and into a destination memory buffer in the cortex, provides the underpinning for symbolic computation. Attractor states linking across the source and destination buffers result in a well-defined symbol vocabulary for such a system. Moreover, we propose that the same thalamic routing operations that subserve such symbolic information relays could also be responsible for orchestrating the training of sub-symbolic cortical networks, and that all of these thalamic operations operate under the control of a discrete action selection system in the basal ganglia and other sub-cortical structures.

Artificial deep neural networks (DNNs) (LeCun et al., 2015) have produced high-performing systems for visual recognition (e.g., (He et al., 2015)), speech recognition and language translation (Amodei et al., 2015)), control (e.g. (Mnih et al., 2015)), and efforts are underway to extend such networks to enable symbolic reasoning (e.g., (Graves et al., 2016; Santoro et al., 2017; Silver et al., 2016)). Increasingly, these artificial DNNs are being considered as a relevant model for aspects of the biological brain’s sensory processing. At the core of these systems are multilayer neural networks based in part on decades-old connectionist models of mammalian cortex (Fukushima, 1980; Hubel and Wiesel, 1962; Rolls and Treves, 1997; Rumelhart et al., 1986b), and trained via a form of optimization that incorporates propagation of credit assignment through multiple layers, e.g., via backpropagation (Rumelhart et al., 1986a). They perform computation in a massively parallel manner that could conceivably map onto cortical anatomy and physiology (Cox and Dean, 2014; Marblestone et al., 2016; Poggio and Anselmi, 2016), and they obey well known biological constraints like the ‘100 step rule’ (Feldman and Ballard, 1982). Recent information theoretic comparisons between artificial DNN vision models and the primate ventral visual stream offer additional, though still preliminary, evidence for the relevance of such connectionist models for understanding biological neural information processing (Yamins and DiCarlo, 2016). Furthermore, the often-cited concerns regarding how backpropagation-based weight training is biologically unrealistic are beginning to be addressed via a set of newer ‘contrastive’ learning rules that are designed to use only information locally available to the synapse, the discovery that requirements on weight symmetry can be relaxed, that the relevant computations can be made asynchronous through synthetic gradients, emerging relations with dendritic compartmentalization and other recent advances (Guerguiev et al., 2017; Jaderberg et al., 2016; Lillicrap et al., 2016; Roelfsema and Holtmaat, 2018; Sacramento et al., 2017). In short, artificial DNNs may capture at least some important aspects of how biological brains function; see (Marblestone et al., 2016) for a recent review.

But a fundamental question remains mostly unaddressed: How should artificial DNNs be viewed in relationship to biological brain architecture as a whole? Should they be viewed as a model of how the entire brain works, or should DNNs instead be viewed as a particular type computational module or sub-component that is utilized within the brain’s larger computational system which itself works along different principles? In this paper we take the later viewpoint, which is, in fact, more along the lines of how DNNs are deployed in today’s AI systems.

The DNNs in today’s AI systems are typically treated essentially as nonlinear classifier modules, or more generally as trainable modules for function approximation, which are embedded in a larger classical ‘symbolic’ computational system that, it can be argued, is doing much of the heavy lifting (Marcus, 2001, 2018; Marcus et al., 2014). For example, this encompassing computational system – in current practice often a software environment such as Python running on a set of von Neumann architecture computers, e.g., (Abadi et al., 2016) – typically ensures that supervised training examples are recorded, stored and presented to the DNN in an appropriate order, provides buffering, control and routing to switch between learning and action, implements the training procedure, records and stores the outputs, and generally provides supporting ‘glue’ or infrastructure – from FOR loops and IF statements all the way up to stochastic gradient descent algorithms – needed to train and utilize the DNN. Although aspects of machine learning aim to remove aspects of the encompassing computational system, potentially allowing future neural network based systems to develop and function more autonomously, the broader question remains as to what structures, if any, might serve analogous “supporting infrastructure” functions with respect to trainable deep networks in the biological brain.

Furthermore, recent “differentiable programming” systems combine trainable connectionist networks with other components such as location-addressable memories (Graves et al., 2016) or tree search algorithms (Silver et al., 2016). Although, in today’s differentiable programming systems, the interaction with these components is often optimized jointly with the connectionist network, e.g., through end-to-end stochastic gradient descent, it remains unclear how the biological brain might orchestrate the interaction of trainable connectionist networks with biologically plausible dedicated memory or routing structures. More generally, how biological neuronal networks might implement rule-based symbolic operations remains a much-debated question, sometimes referred to as the variable binding problem in neural networks (Browne and Sun, 1999; Eliasmith et al., 2012; Hadley, 2009; Hayworth, 2012; Hummel and Holyoak, 1997; Kanerva, 2000; Kriete et al., 2013; Legenstein et al., 2016; Marcus, 2001; Marcus et al., 2014; Plate, 1995; Shastri and Ajjanagadde, 1993; Smolensky, 1990; Touretzky and Hinton, 1985; Zylberberg et al., 2013). Meanwhile, symbolic “production system” models have long been suggested within the cognitive science community as models for the human cognitive architecture (Anderson et al., 2004), and a few attempts at neural implementation of production systems have been made (Eliasmith, 2013; Eliasmith et al., 2012; Lebiere and Anderson, 2008). It nevertheless remains unclear how the brain itself might integrate trainable computation in the style of neural networks, with symbolic computation.

In this paper, we propose how trainable deep learning and classical symbolic computation could co-exist within a single biologically plausible and neuro-anatomically inspired network architecture. We take the view that artificial DNNs trained through some form of multilayer credit assignment may, in fact, comprise a crude first-order conceptual model (Marblestone et al., 2016; Roelfsema and Holtmaat, 2018) for parts of the mammalian cortex, especially for cortical sensory and sensorimotor hierarchies but also for consolidation of higher functions. But crucially, in addition, we suggest that other parts of the cortex, along with subcortical structures, may be better thought of as implementing a more classical, discrete computation and control system, specifically a “production system” (Newell, 1994; Taatgen and Anderson, 2010), consisting of a set of latchable and routable memory buffers supporting symbol transfer, equality detection, and syntactic rule application. In this paper, taking inspiration from the anatomy of cortico-thalamo-cortical relays, we suggest one way that such a production system might be implemented neurally in the brain, and show how it can, in addition, provide the ‘glue’ needed for the brain to train and utilize DNNs.

## Overview of model

This paper describes a set of proposed cortico-thalamo-cortical circuits consisting of:

1. Collections of latching cortical memory buffers, implemented via neural attractor states, which can be trained to share a common ‘symbolic’ vocabulary and which support semantically-consistent copying and comparison operations.
2. Deep neural network hierarchies that can be contrastively (Guerguiev et al., 2017; Hinton and McClelland, 1988; O’Reilly, 1996; Xie and Seung, 2003) supervised-trained to, for example, categorize raw sensory inputs into a memory buffer’s ‘symbolic’ vocabulary.
3. Convergent memory stores able to store and retrieve associations among a set of cortical memory buffers.

Together, these circuits integrate a trainable deep learning system with a classical symbolic system. As we will illustrate, these two linked systems mutually support one another: the deep learning system allows the classical symbolic system to operate on rich sensory data, while the classical symbolic system helps to bootstrap and orchestrate the training of the deep learning system.

As we detail below, the crucial element in each of these circuits will be a model for a set of gateable thalamic higher-order relays. A growing body of anatomical and physiological evidence supports the idea that ‘higher order relays’ in the thalamus provide a means for relaying information among cortical areas (Mitchell et al., 2014; Sherman, 2017). Furthermore, recent evidence has bolstered the view that the persistent storage of information in working memory is mediated by cortico-thalamo-cortical loops, rather than solely via recurrent activation within the cortex itself (Bolkan et al., 2017; Guo et al., 2017). We suggest that the thalamus contains a modest number of these higher-order relays (perhaps on the order of a few hundred) which relay information from one cortical area to another, or back to the same cortical area in order to form a latchable recurrent memory buffer.

By themselves, these cortico-thalamo-cortical circuits would be useless since they all require external control and supervised training. But we are proposing these in the context of a larger theory that assumes that subcortical sources provide such control and training signals, along with recurrent control signals from the cortex itself (Eliasmith et al., 2012; Mitchell et al., 2014; O’Reilly and Frank, 2006; Wei and Wang, 2016). In infancy, control and training signals would derive mainly from subcortical sources like the superior colliculus (SC) and basal ganglia (BG)—structures that are often assumed to rely on self-organized and reinforcement-based learning (e.g., (Doya, 1999; Guillery and Burns, 2017; Mercuri et al., 1997; Redgrave et al., 2010)

These subcortical structures would provide a way to bootstrap the supervised training of DNNs in the cortex which, because of their deep, multi-layered structure and powerful learning mechanisms, will eventually far surpass the computational abilities of the subcortical circuits themselves. After this initial “bootstrapping” phase, we suggest that the cortex itself would learn to control its own thalamo-cortical relays jointly with the subcortical structures.

## Roles for supervised training in the brain

As a brief example of what we mean by ‘bootstrapped’ supervised training, consider a patch of cells along the ventral cortical visual stream in an area destined to become dedicated to representing faces, i.e., a face patch (e.g., (Chang and Tsao, 2017)). Such an area learns to invariantly represent faces, i.e., to respond in the same way to a given person’s face regardless of its orientation or lighting conditions, of whether the person is wearing a hat, of whether the face looks tired or alert, and so forth. How does the brain orchestrate the training of such an area to achieve such invariant recognition?

We suggest that control signals for this process originate from innate face detection circuits in the superior colliculus or other sub-cortical circuits. To train the face patch for invariant recognition, the SC could command the prospective cortical face patch to ‘latch’ or ‘clamp’ in place its current activation pattern when the animal begins attending to a particular, not-yet-view-invariantly-recognizable face. Effectively the prospective face patch would have a “prototype” activation pattern, or “target symbol” pattern, for the novel face latched in place under the control of the SC or other controlling subcortical structure, and kept in place as the child rotates or translates her viewpoint, moves towards the face, interacts with the face, watches its expression change, and so forth.

This latching, along with contrastive Hebbian learning (Ackley et al., 1987; Anderson and Peterson, 1987; Hinton and McClelland, 1988; O’Reilly, 1996; Xie and Seung, 2003) – or with another contrastive learning rule such as (Scellier and Bengio, 2017) or (Guerguiev et al., 2017), or more generally with any learning rule that relies on clamping of neurons at the output layer of a network to provide a supervision signal which is propagated back to earlier layers – would cause the cortical DNN feeding this prospective face patch to learn to associate the various views of the attended face with that particular latched activation pattern. This method of learning view-invariance is really just an extension of the ‘trace rule’ (Földiák, 1991; Wallis et al., 1993) but one that takes advantage of computations and behavioral knowledge available elsewhere in the brain.

Through this method, a simple subcortical face detection circuit might be used to help supervise-train a cortical DNN to perform view-invariant face recognition. Indeed, we will suggest more broadly that this type of cortical latching might be a key mechanism that causes areas of cortex to become specialized, and seek to illustrate how such latching could be neurally implemented and controlled. For example, it is well known that neural responses in higher cortical areas show specializations for object shape, face identity, place, and many other properties (e.g., (Deen et al., 2017; Fedorenko et al., 2011; Saygin et al., 2016; Vul et al., 2012)), and that focal lesions to these areas produce correspondingly specific cognitive deficits (Barton et al., 2002). Our model predicts that some or perhaps all of these specialized cortical areas will be found to have a corresponding thalamic higher-order relay that is innervated by subcortical ‘teaching’ circuits.

Building off of this basic idea, it would be even more advantageous, for DNN training purposes, to allow a more complex teaching circuit to retrieve and latch in an activation pattern recalled from long-term memory when the appropriate conditions arise. For example, if you are reintroduced to a cousin that you have not seen since childhood a (hypothetical) cortical area devoted to representing individuals would likely fall into a completely new pattern of activation due to your cousin’s significant change in appearance over the intervening years. But, because of the verbal reintroduction, ‘you’ know that his now-older appearance should be associated with that previous pattern of activation. In fact it is crucial that this linkage is made in order to allow quick access to previous associations. If part of your brain could recall from long-term memory that previous activation pattern or ‘symbol’ associated with your cousin in childhood and latch it into the appropriate cortical area, then the DNNs of your visual system could be supervised trained to associate his new appearance with that original pattern.

In this paper, we are suggesting that the brain contains just such mechanisms for top-down control of cortical memory recall, routing, and latching operations, and that the cortico-thalamo-cortical circuits we discuss in this paper provide the basic infrastructure for this. In particular, we will suggest that latching or clamping of a cortical memory buffer occurs via a recurrently connected thalamic relay controlled by the basal ganglia via its inhibitory outputs. Moreover, these same latch and relay control operations also, we propose, can underlie a symbolic production system in the brain that may exist in higher-level cortical areas.

## Basal ganglia—a potential source of control signals

This raises the question of what brain structures could provide such control signals—directing (with proper timing) sequences of memory retrievals and routing and latching operations? The reinforcement-trained basal ganglia appears ideally suited to fill this role. The basal ganglia (BG) are an evolutionarily ancient (Grillner et al., 2005; Grillner and Robertson, 2016) set of subcortical nuclei that are thought to form the core action selection circuit in vertebrate brains (Redgrave et al., 1999). BG output consists of a relatively small set of inhibitory “channels” (Grillner and Robertson, 2016; Oorschot, 1996) which provide tonic inhibition to specific populations of cells in the brainstem and thalamus. When a channel’s tonic inhibition is paused, a behavior specific to that population of target cells is ‘released’ (Chevalier and Deniau, 1990; Friend and Kravitz, 2014; Roseberry et al., 2016) The BG does not just release individual actions; it can release precisely-timed sequences of actions (Aldridge and Berridge, 1998; Jin and Costa, 2015), a fact that will be important to our proposal. The BG’s input region, the striatum, is a shallow neural network (Friend and Kravitz, 2014; Reynolds and Wickens, 2002) whose synapses undergo LTP/LTD modulated by dopamine signals from the brain’s reward system, e.g., the ventral tegmental area (Kreitzer and Malenka, 2008; Yagishita et al., 2014). The brain’s reward system is modulated by primary reward signals as well as by the striatum itself and by other areas, creating a combined system optimized for temporal difference reinforcement learning and other forms of reinforcement learning (Hazy et al., 2007; O’Reilly et al., 2007, 2014). Thus, in our model, as in much past work (e.g., (Eliasmith, 2013; O’Reilly and Munakata, 2000)), the BG comprises a reinforcement-trained action selection system, based on shallow pattern recognition, for which selecting an action corresponds to disinhibition of a downstream circuit.

Most importantly for our proposal, a subset of thalamic relays with projections to and from cortical areas involved in sensory, motor, memory, emotional, and cognitive functions appear to be under the direct inhibitory control of the basal ganglia (Clower et al., 2005; McFarland and Haber, 2002; Middleton and Strick, 1996). The basal ganglia projections to higher-order thalamic relays associated with visual areas in the parietal cortex (Clower et al., 2005) and inferotemporal cortex (Middleton and Strick, 1996) are particularly noteworthy in light of our proposal as these projections could provide the type of latching and routing signals we suggest are used to train the many specialized areas found in these visual cortical regions. Thus, basal ganglia inhibition/disinhibition could gate thalamic relays on and off, thereby controlling information storage and transmission among cortical memory buffers.

In summary, the BG seems anatomically capable of providing the types of timing and control signals assumed by our proposal to orchestrate latch and relay control, via its inhibitory channels to higher-order thalamus. Interestingly, studies have shown that when an animal’s neocortex is removed early in life while sparing striatal circuits, the animal retains significant capacity for intelligent behavior, including learning, much of which can likely be traced to the spared action selection circuits of the BG (e.g., (Sorenson and Ellison, 1970)). Quoting from Grillner et al. (2005): “[Animals] decorticated at birth move around gracefully and display goal-directed locomotion – the uninformed observer would see no or little difference in movement pattern from that of an animal with its cortex intact.”

We do not directly model BG or SC computations in this paper. Instead, we focus solely on modeling the aforementioned classes of cortico-thalamo-cortical circuits – latching memory buffers, transfer of information through thalamic relays, supervised-trained DNNs, and convergent memory stores. But in the process of modeling these, we will be asserting that their thalamic relays are being controlled by precisely-timed sequences of inhibition, and we are assuming that such control sequences derive from subcortical structures like the BG.

## Symbolic computations enabled by the architecture

We have so far discussed the idea of supervise-training a cortical DNN by latching a particular pattern of activation into its ‘last’ layer—a cortical buffer meant to model a specialized region high in the cortical hierarchy. In a sense, each specialized cortical buffer can be assumed to have a ‘symbol’ vocabulary consisting of all the stable patterns of activation that that buffer supports, each symbol being semantically grounded in the real world through the action of its attached sensory DNN. But a collection of such neural buffers cannot, by themselves, support true symbolic computations because they cannot support symbol binding (Hadley, 2009; Marcus, 2001), since there is no natural way to “translate” or “copy and paste” arbitrary contents of one buffer into another buffer – after all, each buffer uses a different, learned encoding scheme, so there is no way, by default, to compare their contents. This is the classic neural binding problem, where we have defined variable binding (Marcus et al., 2014) as “the transitory … tying together of two [pieces] of information: a variable (such as an X or Y in algebra, or a placeholder like subject or verb in a sentence) and an arbitrary instantiation of that variable (say, a single number, symbol, vector, or word)”. What is needed, but not solved by a naïve system of cortical memory buffers, is a way to assign arbitrary “filler” content to an arbitrary “role”, “slot” or “variable”.

To overcome this limitation, our models assume that every sensory DNN ends in a collection of cross-connected cortical buffers which together form a Dynamically Partitionable Auto-Associative Network (DPAAN) (Hayworth, 2012). All of the buffers in a single DPAAN are trained simultaneously to share a *common set of stable attractor states*, each state corresponding to one symbol. We show that the collection of buffers in a DPAAN can support symbol binding by allowing semantically consistent buffer-to-buffer copying and equality detection of symbols, i.e., of a vocabulary of stable activity patterns. Buffer-to-buffer equality detection is crucial to allowing a shallow pattern recognizer network, like that in the striatum, to learn syntax-based rules (Hayworth, 2012), and buffer-to-buffer copying is crucial to allow the learning of rules that manipulate syntax. The nature of these operations will be discussed in more depth in the **Discussion** section, but it is important to understand that they are key to our assertion that the cortico-thalamo-cortical circuits we describe in this paper are sufficient to support production system-style computations (Taatgen and Anderson, 2010).

## Relations with other proposals

Our paper attempts a novel synthesis of ideas existing in the literature. For example, our mathematical model of attractor based working memory derives from bidirectional associative memory (Kosko, 1988) and from classic models of hippocampal memory storage (Lisman et al., 2005; Rennó-Costa et al., 2014; Rolls, 2010). Our proposal that working memory is jointly stored between thalamus and cortex owes much to the FROST model (Ashby et al., 2005). Our picture of thalamic relays is inspired by the work of Sherman and Guillery as well as others on the anatomy of thalamo-cortical connections (McFarland and Haber, 2002; Sherman, 2017). Our suggestion that the basal ganglia provides control over thalamo-cortical memory structures via inhibition and disinhibition of the thalamus is similar to that of O’Reilly and Frank (O’Reilly and Frank, 2006), and of (Stocco et al., 2010), as well as (Stewart et al., 2010). This view of the basal ganglia output to the thalamus is supported by recent experimental studies in songbirds (Goldberg et al., 2013). More broadly, the view of the basal ganglia as a discrete action selection system derives from (Redgrave et al., 1999) and is also used in the ACT-R literature based on the idea of the basal ganglia as controlling a production system (Stocco et al., 2010). The suggestion that the basal ganglia orchestrates training of cortical networks derives from (Ashby et al., 2010) and was revisited in (Pyle and Rosenbaum, 2018), and the notion that hippocampal information is consolidated into cortex for permanent storage is prevalent in works such as (Atallah et al., 2004; Kumaran et al., 2016; McClelland et al., 1995). Finally, the idea of contrastive training of neural networks, and of providing supervision signals to a network by clamping or perturbing an output layer has been important in the machine learning and computational cognitive science fields for some time (e.g., (O’Reilly, 1996; Xie and Seung, 2003)) and continues to be today (e.g., (Guerguiev et al., 2017; Scellier and Bengio, 2017)). Our overall architecture linking deep hierarchies, symbolic intermediates supporting discrete operations, and reinforcement learning, is reminiscent of (Garnelo et al., 2016). See the **Discussion** section for many other connections with the literature, although unfortunately we do not have space here for a comprehensive review of the relevant precursors.

## Results

In what follows, we first introduce and simulate a connectionist model of a controllable working memory buffer in which information is stored via persistent activity in a cortico-thalamo-cortical loop (**Simulation #1**). The dynamics of this loop creates a finite set of attractor states, where each attractor state is proposed to correspond to one ‘symbol’ of the buffer’s vocabulary. This model builds on other similar models of working memory (Ashby et al., 2005; O’Reilly and Frank, 2006).

We then simulate the transfer of symbols between multiple such buffers, via relays, and the comparison (equality detection) of contents across multiple buffers (**Simulation #2**). This controllable working memory model serves as a more biologically plausible implementation of the Dynamically Partitionable Auto-Associative Network (DPAAN) concept originally described in (Hayworth, 2012), which comprises a conceptual solution for symbolic computation and “variable binding” (Hadley, 2009; Kriete et al., 2013; Legenstein et al., 2016; Marcus, 2001; Marcus et al., 2014) in a neural substrate. DPAAN is an example of an “anatomical binding” solution to the variable binding problem, wherein each variable corresponds to an anatomically separate register. Such solutions face a general problem of ensuring that symbols can be compared and transferred across buffers, despite different coding schemes used in each buffer. DPAAN enables this via a large-scale network linking many buffers. A global symbol comprises an attractor state of this network. In the original DPAAN formulation, synapses between buffers within the global network could be gated on and off, thus linking attractors in each buffer into a single attractor of the global network, or separating buffers such that each one can take on its own attractor state without interference from the attractor states occupied by the other buffers. (Hayworth, 2012) showed how such a scheme can allow the “copying and pasting” of symbols between buffers, as the state of a source buffer forces a destination buffer into a corresponding state that is consistent at the global level, when the buffer-to-buffer synapses are gated on. The model presented here comprises an alternative implementation of DPAAN in which thalamus-inspired latched buffers and relays, rather than directly controllable synapses, enable the buffer to buffer transfer operations.

Next, we exploit these latched buffers and controllable relays to illustrate the training of a multi-layer feedforward neural network using a contrastive Hebbian learning scheme (**Simulation #3**). In effect, this demonstrates how the symbols in a DPAAN can be grounded to sensory percepts.

Finally we combine these ideas with a simple hippocampus-inspired declarative memory model (**Simulation #4**). The goal of this model is to demonstrate how multiple DPAANs can be used to properly bind the features of a multi-object scene, and, importantly, to demonstrate how this entire multi-object scene representation can be ‘flattened’ and stored as synaptic changes in an attractor-based declarative memory. Further, we demonstrate how controlled routing allows for the associative recall of specific elements of a stored multi-object scene.

The connectionist models described are purposely simplified (e.g., binary activations, k-winners-take-all dynamics, discrete time steps) as they are meant only to test and demonstrate the key principles we are describing in this paper. In the discussion, we will discuss the limitations of this model, detail what is known and unknown about its potential neuroanatomical underpinnings, relate the model to other work in production system models and machine learning, and suggest future experimental and computational directions.

### Simulation #1: Cortical buffer-to-thalamic latch bidirectional associative memory

At the core of each DPAAN module is a set of cortical memory buffers each of which can hold one symbol. Each cortical memory buffer has an associated thalamic latch to which it is bidirectionally connected (Figure 1). The pair forms a bidirectional associative memory (BAM) (Kosko, 1988) but implemented with k-winners-take-all (k-WTA) dynamics. If it is assumed that the thalamic half of this BAM is under the inhibitory control of, for example, the basal ganglia, then this provides a simple means to “latch” a memory into the BAM at a desired time. When the thalamic part is uninhibited, the cortical and thalamic parts recurrently excite one another, whereas when it is inhibited, the thalamic part has zero activation (inhibition operates by setting the k-WTA parameter, k, to zero in the latch) leaving the cortical part to fall into a state dictated by internal noise and external input. This arrangement has been used in other models of short term memory (e.g., (Ashby et al., 2005; O’Reilly and Frank, 2006)) and has recently received some experimental support in a study of working memory in the mouse in which the thalamic part of a presumed cortico-thalamo-cortical memory loop was silenced optogenetically (Guo et al., 2017).

**Figure 1.**
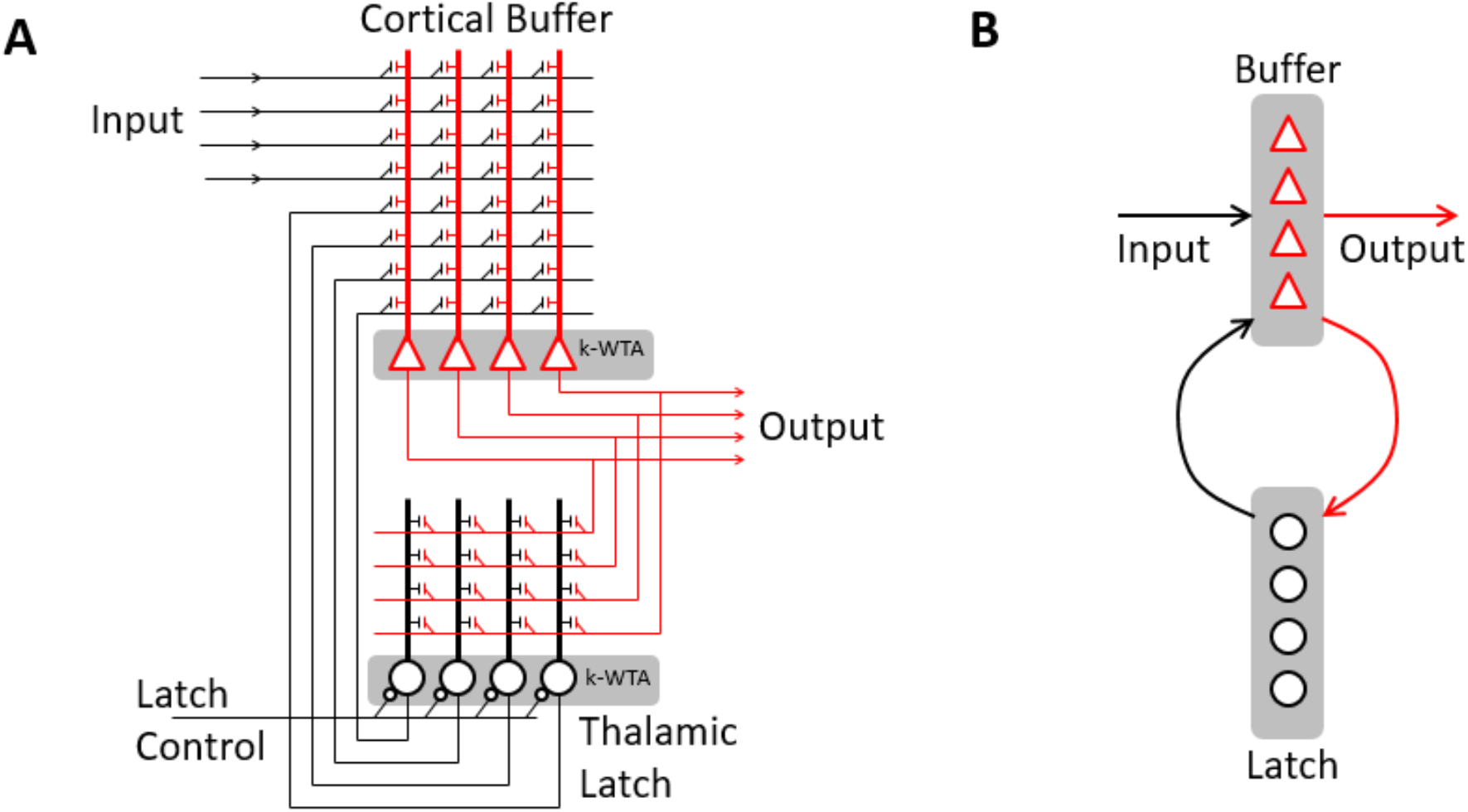
Thalamocortical latching memory buffer. (A) Each cortical memory buffer has an associated thalamic latch to which it is bidirectionally connected creating a bidirectional associative memory (BAM). Both the cortical buffer and the thalamic latch use k-winners-take-all (k-WTA) dynamics to ensure a specified sparseness of activation. This diagram shows the synaptic weight matrices connecting the two which are trained to give the BAM a set of stable attractor states. When the thalamic latch is inhibited the cortical buffer is solely driven by external input (Input). When the thalamic latch is later disinhibited, the BAM will fall into the attractor state most closely associated with the input and will remain there even when the input is removed or changed. (B) Simplified diagram (used later) where each synaptic matrix is depicted as a single arrow connection.

To test this arrangement, we first modeled a single cortical buffer-to-thalamic latch memory. The cortical buffer consisted of *n* = 300 neurons and its corresponding thalamic latch consisted of *p* = 300 neurons (note, these do not have to be equal). These were connected via weight matrices ***W***^buffer to latch^ and ***W***^latch to buffer^ initialized with weights sampled independently from a uniform distribution spanning 0.0 to 0.2. A set of *L* = 100 randomly generated ‘memories’ was created to train the network; these memories will be the symbols in the full DPAAN architecture. Each symbol *S_l_* (*l* ∈ {1 … *L*}) consisted of a pair of randomly generated sparse activation patterns 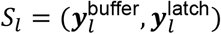 where 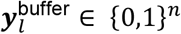 and 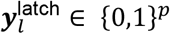. The sparseness (*α*) of these vectors was set to be 0.05, corresponding to *k*_buffer_ = *n* ∙ *α* and *k*_latch_ = *p* ∙ *α* active neurons for the k-WTA dynamics of the buffer and latch respectively.

Training of the weight matrices was as follows. The activation pattern of each unique symbol *S_l_* to be learned was put into the buffer and latch, and Hebbian weight matrix increments were computed as follows (vectors ***y***^buffer^ and ***y***^latch^ are assumed to be column vectors):

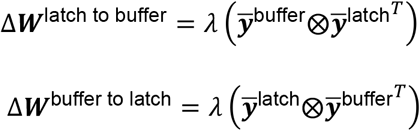

Where ⊗ is the outer product and *λ* is a learning rate parameter which was set to 1.0 in this simulation. The vectors 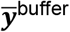 and 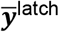 were computed to be zero mean as follows:

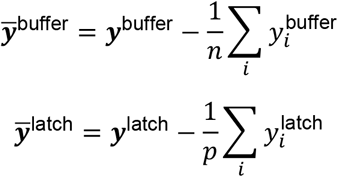

This ensured that the delta weight matrices summed to zero overall ensuring that weights would remain bounded.

Each symbol *S_l_* was trained in order 1 … *L* with the update rules:

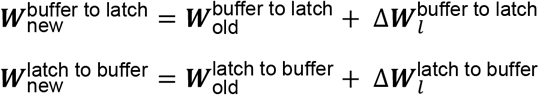

After each weight matrix updated, any weight greater than one or less than zero was set to 1.0 and 0.0 respectively. Use of these update rules meant that more recent symbols would slowly overwrite older ones even in cases where the sparse patterns did not overlap.

After training, recall of each symbol was tested by initializing the buffer’s activation pattern with a degraded version of that symbol’s 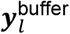 pattern along with a low level of random activation noise. A 40% degraded pattern had 0.6 · *k* of the original *k* active neurons set correctly, so that these would definitely be in the k-WTA group along with a randomly selected additional 0.4 · *k* neurons.

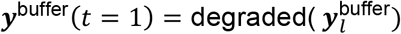

Then the network was simulated in discrete time steps in the following order:

**Sub-step #1**: Update latch activation ***y***^latch^:

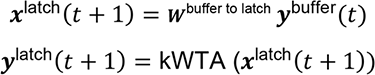
**Sub-step #2**: Update buffer activation ***y***^buffer^:

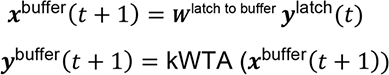

This sequence was repeated for 10 time steps and the resulting pattern of firing in the buffer was compared against the target pattern (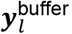) that should have been recalled. Recall was deemed successful if the final pattern had greater than 90% correlation with that target. The results of several such tests of this latching memory are summarized in the **Supplementary Online Material**. As long as the number of symbols trained is kept below the network’s maximum capacity then memory recall is perfect and the network provides ‘clean up’ for significantly degraded initial patterns. (See (Kosko, 1988) for a more complete treatment of the general capacity of BAM memories.) If the network is trained beyond its memory capacity, then more recent symbols overwrite older ones. As noted above, this is due to our clipping of synaptic weights after each memory is trained. This was the behavior we desired for our memory buffer since we theorize that the brain will choose to generate a new DPAAN symbol on demand (by generating a random activation pattern) when presented with a truly novel world feature it must represent.

When the thalamic latch neurons are inhibited, by programmatically setting their k-WTA target to k=0, the reverberating BAM’s memory is lost. This implementation of inhibition constitutes a simplified model (O’Reilly and Munakata, 2000), but similar effects are expected for more realistic implementations of global inhibitory inputs to a buffer or latch population.

### Simulation #2: Basic DPAAN functionality – buffer-to-buffer symbol transfer and equality detection

Figure 2 depicts a basic DPAAN network consisting of two latchable buffers interconnected by switchable thalamic relays and an equality detection network. The key to the DPAAN’s functionality is the fact that it is trained as a single interconnected autoassociative network. The network is trained as a whole such that global patterns of activation (designated *S_l_* below) become stable attractor states of the entire network; each attractor state corresponding to one symbol in the DPAAN’s vocabulary. In this way, disinhibiting, for example, ***y***^relay1→2^ reconstitutes part of the original global network forcing ***y***^buffer2^ to fall into the same attractor state that ***y***^buffer1^ is in.

**Figure 2.**
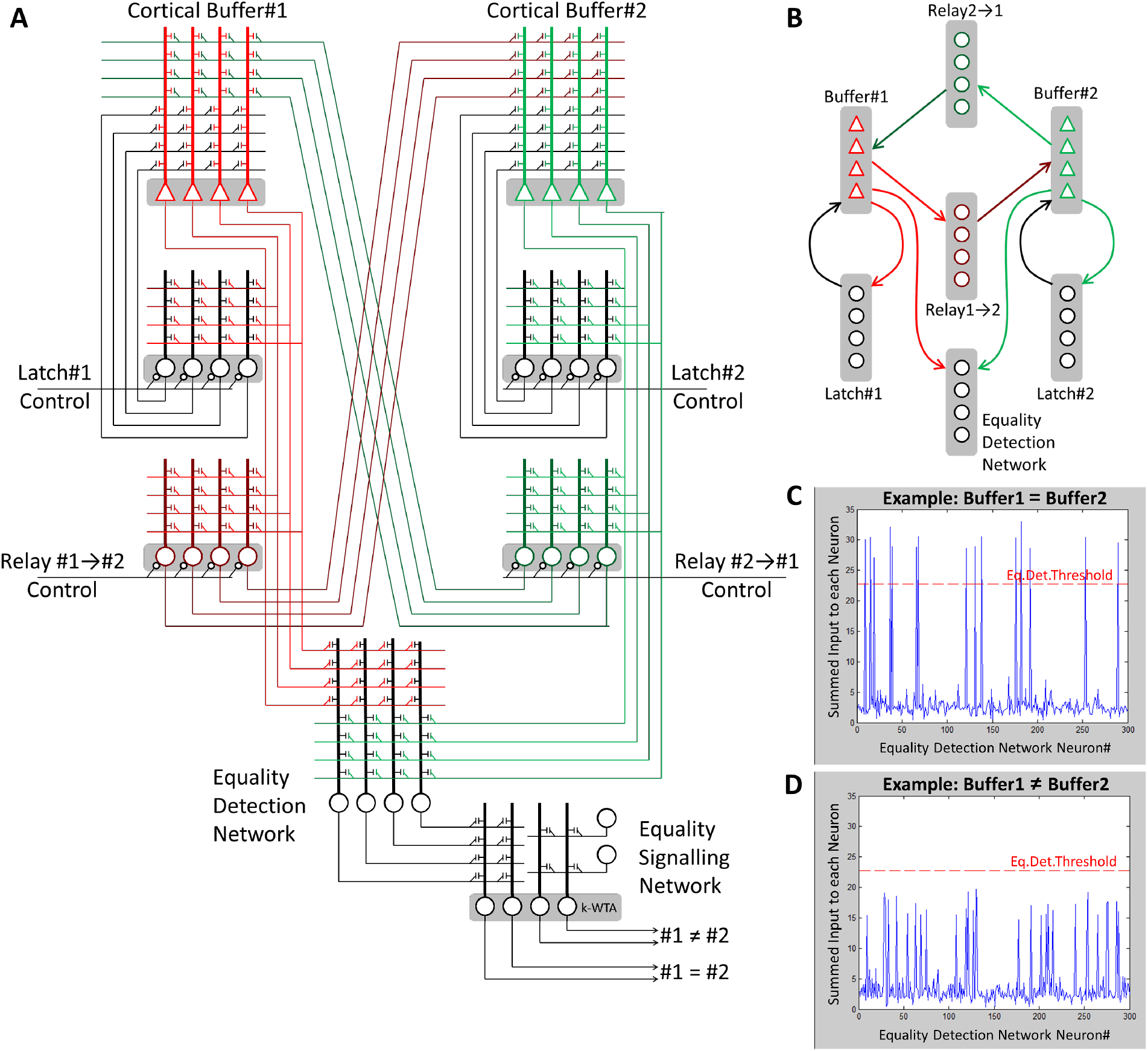
Most basic DPAAN network – two latchable buffers interconnected by transfer relays and an equality detection network. (A) A single two-buffer DPAAN network consisting of seven neural populations (***y***^buffer1^, ***y***^latch1^, ***y***^relay1→2^, ***y***^buffer2^, ***y***^latch2^, ***y***^relay2→1^, ***y***^Eq.Det.Net^). All of the weight matrices are trained simultaneously so that the full network will have a set of global attractor states, these attractor states forming the symbol vocabulary of the DPAAN. Inhibiting the relay networks ‘partitions’ this global network into two independent BAM memories. Disinhibiting Relay#1→#2, for example, allows Buffer#1 to drive Buffer#2, thus implementing a buffer-to-buffer symbol copy operation. (B) Simplified diagram where each synaptic matrix is depicted as a single arrow connection (used later). (C,D) Plots showing input current levels in the equality detection network prior to thresholding. When buffer #1 and #2 are storing the same symbol more neurons cross this threshold. In this way the network is continuously signaling whether buffers #1 and #2 are representing the same symbol.

We modeled such a DPAAN network using Matlab. Each neuronal group (buffers, latches, relays, eq. det. network) consisted of *n* = 300 neurons. These were connected via weight matrices ***W***^buffer1 to latch1^, ***W***^buffer1 to relay1→2^, and so forth, initialized with weights sampled independently from a uniform distribution spanning 0.0 to 0.2. A set of *L* = 100 randomly generated symbol ‘memories’ were created to train the network. Each symbol *S_l_* (*l* ∈ {1 … *L*}) consisted of a set of randomly generated sparse activation patterns 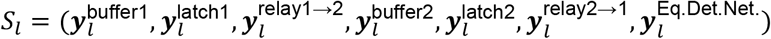, where 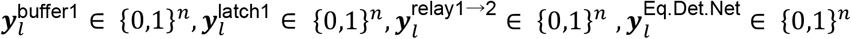, and so forth.

Training of the weight matrices was performed similar to the single buffer case above, using Hebbian association. The activation pattern of each unique symbol *S_l_* to be learned was loaded into all neuronal groups, and delta weight matrices were computed as follows:

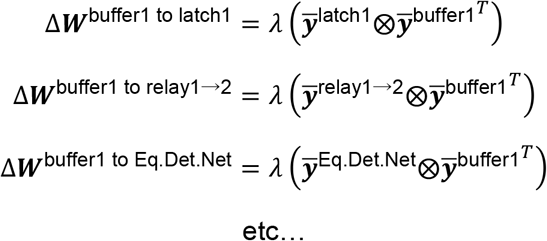

As previously, these deltas were used to update their corresponding weight matrices and the updated weights were clipped to fall in the range 0.0 to 1.0 after each memory was trained.

The buffers, latches, and relays all use k-WTA dynamics during simulation, but the equality detection network does not. Instead, firing in the equality detection network is determined by comparing each neuron’s net input to a threshold:

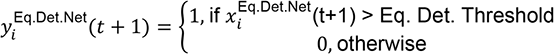

Recall that weight inputs to the Eq. Det. Network are trained along with all other parts of the full DPAAN as part of a global autoassociative network. If Buffer#1 and Buffer#2 are currently storing the same symbol (e.g., patterns 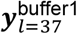 and 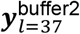 corresponding to global attractor state *l* = 37) then those Eq. Det. Network neurons associated with that trained symbol (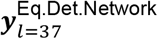) will receive maximum net input current; in particular, there will be k = *n* · *α* neurons in the equality detection network which receive high input current, with the rest receiving low input current (Figure 2C). However, if Buffer#1 and Buffer#2 are currently storing *different* symbols (for example 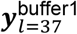 and 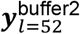) then the resulting input current pattern ***x***^Eq.Det.Network^ will be more spread out across neurons, with typically 2k=2 *n* · *α* neurons receiving an intermediate level of input current (Figure 2D). In this case, there will be more driven neurons, but their input currents will be less than if Buffer#1 and Buffer#2 were currently storing the same symbol and thus the driven neurons will not reach the threshold level of input current necessary for activation. It is this difference that we exploit to determine equality. The value of Eq. Det Threshold is chosen to ensure this behavior.

We determine the ‘optimal’ value of the Eq.Det.Threshold parameter by loading in 50 example cases where buffer1 and buffer2 contain the same symbol and determine what average threshold would be necessary in each of those cases to allow k neurons to fire. Then we load in 50 example cases where Buffer#1 and Buffer#2 contain *different* symbols and determine what average threshold would be necessary to allow k neurons to fire. The Eq.Det.Threshold parameter is set to be half way between these two values. During normal operation, this means that typically k neurons in the Eq. Det. Network will fire when the two buffers contain the same symbol and *none* will fire when they contain different symbols. Examples of this are displayed in Figure 2.

An Equality Signaling Network is used to convert this k or none output into one in which half of the neurons fire to signal ‘equal’ and the other half fire to signal ‘not equal’. This is more suitable for training downstream pattern association networks, for example circuits in the striatum. Recall that the whole point of training a set of equality detection networks is to allow simple downstream decision making circuits to learn syntax-based rules that are sensitive to the bound contents of variables, for example: ‘if Var1 == Var2 then …’. The Equality Signaling Network is set up to make the comparison of variable contents sufficiently explicit that such simple pattern matching ‘rules’ in the striatum could potentially learn (via reinforcement) to be triggered by such equality. (Hayworth, 2012) provides a more in depth discussion of these ideas.

For testing, a three buffer DPAAN was simulated. The three buffers and latch activation vectors were initialized with 40% degraded patterns corresponding to three different symbols chosen at random. The goal was to let these settle into symbol states in each buffer, then to disinhibit a specified relay to copy the symbol of one of the buffers (the ‘source’ buffer) into another (the ‘target’ buffer), overwriting its previous contents. Success of this operation was evaluated by comparing the final buffer contents with the desired patterns, and by checking that all equality networks correctly signaled which buffers were equal and which were not equal.

At the start of each test, all latches were initialized as disinhibited so that these degraded patterns would be cleaned up and stabilized. All relays were inhibited initially so that each buffer-latch memory acted independently. The network was then simulated in discrete time steps in the following order:

**Sub-step #1**: Update buffer activations
**Sub-step #2**: Update latch activations
**Sub-step #3**: Update relay activations
**Sub-step #4**: Update eq. det. network activations

This cycle was repeated for three time steps to allow initial settling of the buffer-latches. Then on the 4^th^ time step the latch associated with the target buffer was inhibited, eliminating the reverberating activity that kept that buffer’s pattern stored, preparing it to accept new input. On the 5^th^ time step the relay from source to target was disinhibited. This reconstituted the part of the original autoassociative network connecting the source buffer to the target buffer, allowing the source buffer to drive the target buffer into its attractor state. On the 6^th^ time step the target buffer’s latch was disinhibited, locking the new contents in place. On the 7^th^ time step the relay from source to target was inhibited, again rendering all three buffers independent. The simulation then continued until time step 10 when the final patterns were evaluated. This buffer-to-buffer transfer sequence is depicted in Figure 3.

**Figure 3.**
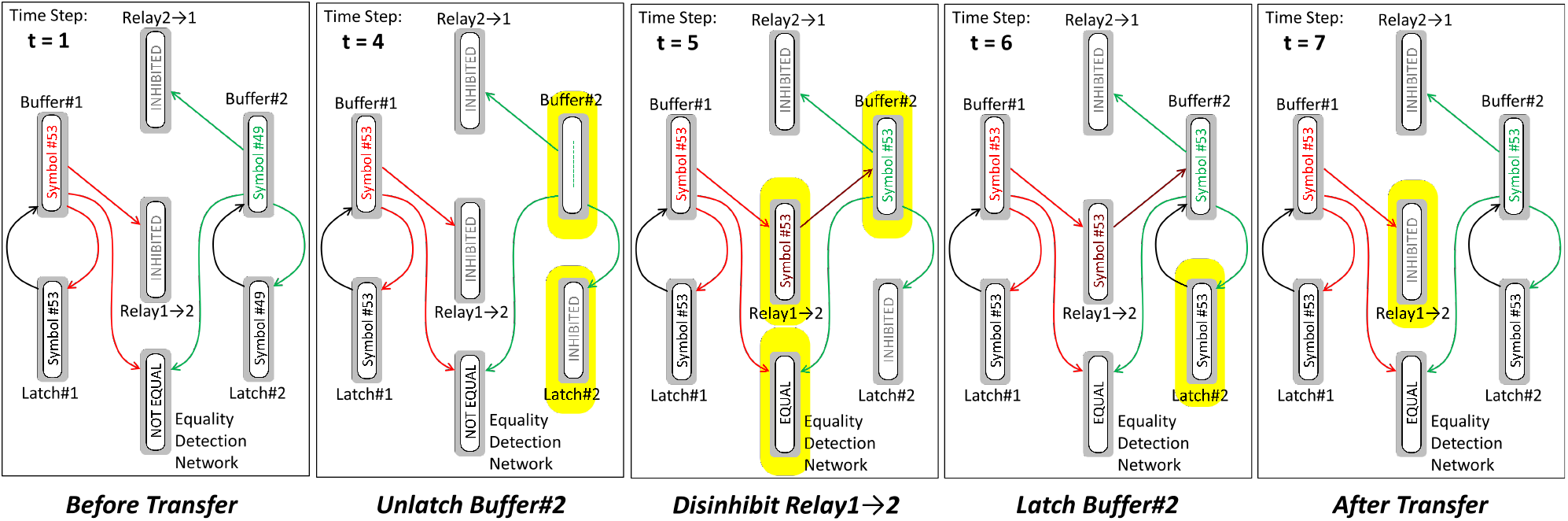
Diagram showing the steps of a buffer-to-buffer transfer. In this case Symbol#53 is copied from Buffer#1 to Buffer#2. Neuronal groups that change during each time step are highlighted in yellow.

In our simulations we found that as long as the number of symbols trained is kept below the individual BAM memories’ maximum capacity then such copy and equality detection operations are performed flawlessly.

### Simulation #3: Contrastive Hebbian learning (CHL) using DPAAN buffer transfers and latching

As we have seen, a single DPAAN module consists of a set of memory buffers each of which shares a common finite vocabulary of symbols. But how are these symbols grounded in the real world? We hypothesize that such ‘symbol grounding’ is handled by a supervised-trainable input network which provides an interface between the earlier stages of an agent’s sensory processing network and the DPAAN module’s first buffer. Figure 4 depicts a DPAAN module augmented with such an input network consisting of two ‘hidden’ layers. The job of this input network is to act as a classifier network mapping the presented sensory input to one of a discrete set of categories (i.e., one of the DPAAN’s symbols).

**Figure 4.**
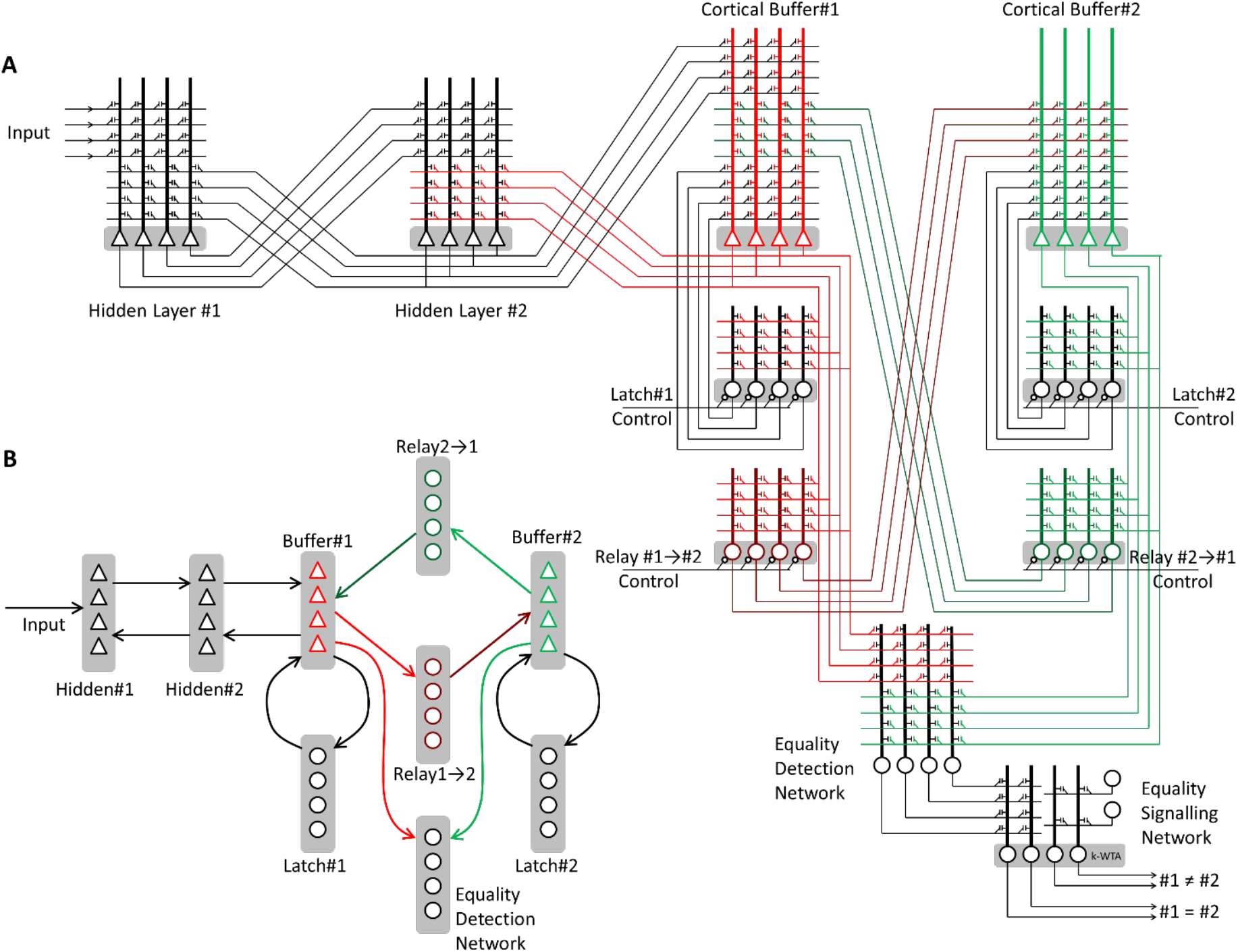
DPAAN network augmented with a multilayer input network. (A) Diagram depicting a single two-buffer DPAAN network like that shown in Figure 2 but modified to include a deep input network providing input to Buffer#1. Contrastive Hebbian learning is used to train this deep input network to associate DPAAN symbols with particular input categories. (B) Simplified diagram where each synaptic matrix is depicted as a single arrow connection (used later).

Supervised training of deep classifier networks (by, for example, backpropagation) is commonplace, but it is an open question how such supervised training might occur in the biological brain. Here we suggest that this training occurs via some form of contrastive learning with multi-layer credit assignment, e.g., Contrastive Hebbian learning (Xie and Seung, 2003), where the two phases of CHL are controlled by the DPAAN’s relays.

The ‘minus’ and ‘plus’ phases of this CHL training procedure are depicted in Figure 5. The target pattern to train toward is first latched into DPAAN Buffer#2. If ***y***^latch1^ and ***y***^relay2→1^ are both inhibited then DPAAN Buffer#1 will only be driven by the input network (i.e., by hidden layer #2). This represents the ‘minus’ CHL phase where the network’s activation is driven solely by the sensory input. The input network has both feedforward and feedback connections, so during the ‘minus’ CHL phase all hidden layers will come to an equilibrium state of activation based solely on input driving. This activation pattern is called the ‘minus’ phase equilibrium state and is recorded for later use. Following this, ***y***^relay2→1^ is disinhibited thus initiating the ‘plus’ CHL phase where the desired target pattern is clamped. In our simulations we make the synaptic drive from the ***y***^relay2→1^ onto Buffer#1 many times greater than the drive from hidden layer #2 onto Buffer#1, which (along with the k-WTA dynamics) effectively clamps the target pattern into Buffer#1. The recurrent connections then cause the hidden layers to fall into a new activation pattern called the ‘plus’ phase equilibrium state. The hidden layers’ weight matrices are then updated according to the CHL learning rule (Xie and Seung 2003) based on these recorded ‘minus’ and ‘plus’ phase equilibrium states.

**Figure 5.**
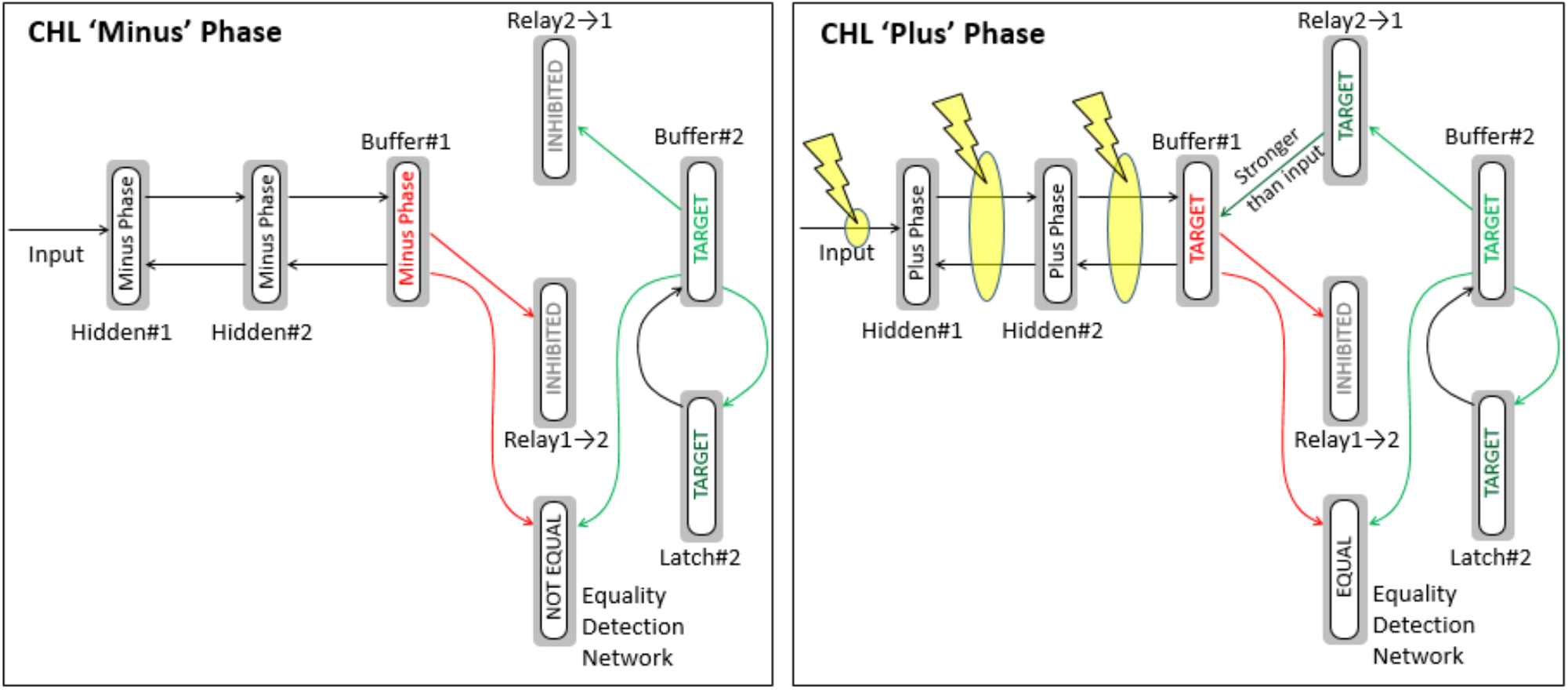
Training the deep input network by Contrastive Hebbian Learning. ‘Lightning bolt’ symbols indicate those weight matrices that are trained by the CHL rule.

#### Details of simulation

In order to test this idea we created a set of 70 letter images. These consisted of 10 letter shapes (A,F,I,J,N,Q,S,V,W,X) each depicted in 7 different colors (white, red, green, blue, yellow, magenta, cyan) on a black background (28 × 28 pixels). The RGB pixel images of these letters were flattened to produce vectors with a length of 2352 values (28 × 28 × 3). These 70 input vectors were used to train the network. For this simulation two DPAAN’s were used, each with its own input network. DPAAN#1 was trained to classify based on letter shape and DPAAN#2 was trained to classify based on color. This was done to help demonstrate the idea that multiple DPAANs, each trained separately to represent different aspects of the same object, can be used together to solve the neural binding problem as depicted in Figure 6.

**Figure 6.**
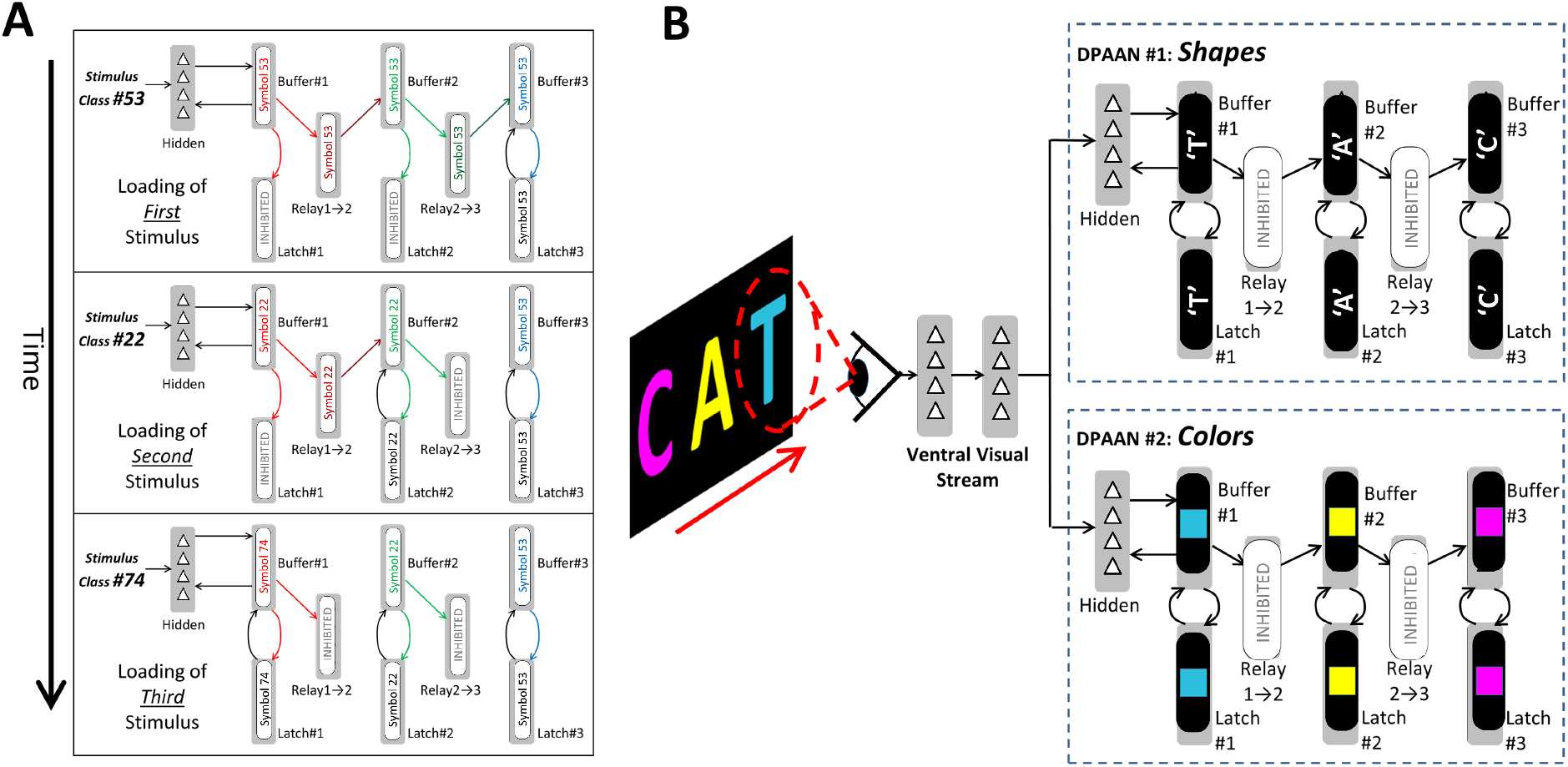
Sequential loading of DPAANs. (A) Steps used to convert a temporal sequence of inputs into a ‘flat’ pattern of activation across DPAAN buffers. This is accomplished by sequentially inhibiting DPAAN relays. The first stimulus (Class#53) is initially represented simultaneously across all DPAAN buffers as Symbol#53. A later presentation of a second stimulus (Class#22) causes Symbol#22 to overwrite all but Buffer#3. Finally presentation of a third stimulus (Class#74) causes overwriting of Buffer#1 with Symbol#74. The result is that this three stimulus temporal sequence has been encoded by a ‘flat’ representation where each of the three symbols is assigned to a unique cortical buffer. (B) Example of how multiple DPAANs can be used to represent different aspects of the same viewed object (in this case shape and color), and how multiple DPAANs can together create a structured, correctly bound representation of a multi-object scene.

Ten of DPAAN#1’s internal symbols were used, one to represent each of the ten letter shape categories. Similarly, seven of DPAAN#2’s internal symbols were used, one to represent each of the seven color categories. We here describe the procedure used to train DPAAN#1 on letter shape classification. The training of DPAAN#2 was similar but for color classification.

DPAAN#1 was setup as described in the previous section to have three buffers with their associated latches, relays, and equality detection networks, and to be able to represent a total of 20 symbols. Ten DPAAN#1 symbols were chosen to represent the ten letter shape categories. The input network consisted of an input layer with 2352 neurons, and two hidden layers each having 900 neurons. Each hidden layer obeyed k-WTA dynamics with a sparseness of 0.05. As depicted in Figure 4, there are five weight matrices to train in the input network: ***W***^input to hidden1^, ***W***^hidden1 to hidden2^, ***W***^hidden2 to buffer1^, ***W***^buffer1 to hidden2^, ***W***^hidden2 to hidden1^. The first three of these represent feedforward connections while the latter two are feedback connections. Their weights were all set to random values prior to training.

Training consisted of 20 epochs in which all 70 example input pictures were presented in randomly permuted order. The input layer’s activation ***y***^input^ was set to an example picture’s flattened vector representation, and the desired target symbol was latched into DPAAN#1’s ***y***^buffer2^. ***y***^latch1^ and ***y***^relay2→1^ were both initially inhibited so that ***y***^buffer1^ was driven only by the input network via connections from ***y***^hidden2^.

The network was then simulated in discrete time steps in the following order:

**Sub-step #1**: Update hidden1 layer activation
**Sub-step #2**: Update hidden2 layer activation
**Sub-step #3**: Update buffer activations
**Sub-step #4**: Update latch activations
**Sub-step #5**: Update relay activations
**Sub-step #6**: Update eq. det. network activations

This cycle was repeated for four time steps to allow the ***y***^hidden1^, ***y***^hidden2^, and ***y***^buffer1^ to settle into a stable configuration. These firing patterns were then recorded as 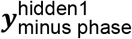, 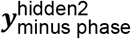, and 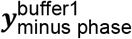. At time step #5, ***y***^relay2→1^ was disinhibited allowing buffer#2 to drive ***y***^buffer1^ into the ‘target’ symbol pattern. The simulation was run for another five time steps and the resulting firing patterns were recorded as 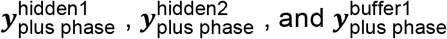. Updates to the five weight matrices are then computed as follows:

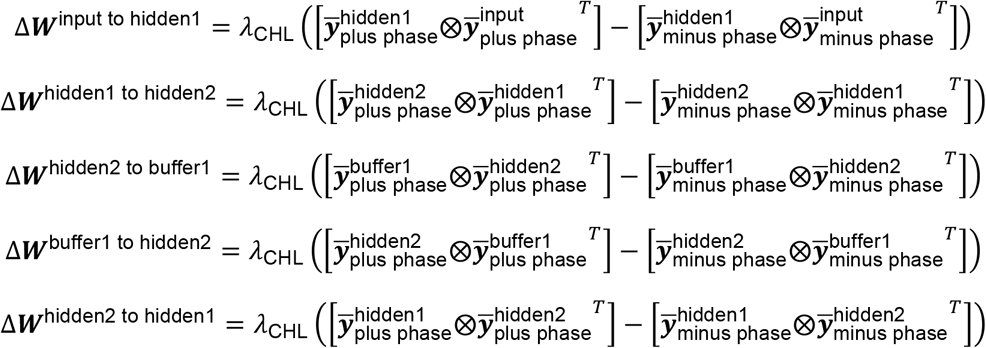

A learning rate of *λ*_CHL_ = 0.000003 was used. At the end of 20 training epochs, DPAAN#1 was able to correctly classify all of the 70 example images based on letter shape, and DPAAN#2 was able to correctly classify all of the 70 example images based on their color.

We used colored letter stimuli for testing to demonstrate how two separate DPAANs can each be trained to represent one aspect of a sensory input, i.e. DPAAN#1 representing shape and DPAAN#2 representing color. However judging the difficult of the classification task is not easy using such stimuli, so we ran an additional challenging classification task which sought to classify a collection of 225 random overlapping binary vectors into 15 different categories.

We generated a set of 225 random input vectors 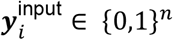 of length *n* = 1000 and sparseness 0.2. Each of these vectors was randomly assigned to one of 15 ‘shape’ classes and one of 15 ‘color’ classes to be learned by DPAAN#1 and DPAAN#2 respectively. Category assignments were generated so that all (shape X color) combinations were represented in the 225 vectors. For this task each DPAAN was provided with an input network containing three hidden layers, each layer having 900 neurons and a k-WTA sparseness of 0.05. After a total of 11 CHL training epochs the deep DPAAN input networks had learned to perform this categorization task perfectly. A summary of these results is included in the **Supplementary Online Material**.

### Simulation #4: Declarative memory network for two DPAAN modules sharing a common input

As shown in the previous section, a system containing multiple DPAANs can be constructed so that each DPAAN represents a distinct aspect of the sensory input. DPAAN#1’s input network was trained to classify the sensory input based on shape, and DPAAN#2’s input network was trained to classify the sensory input based on color. Figure 6B shows how such a multi-DPAAN system might be used to represent a multi-object scene as it is sequentially attended. First the magenta letter ‘C’ is attended allowing its shape to be classified and stored in DPAAN#1’s 3^rd^ buffer, and its color to be classified and stored in DPAAN#2’s 3^rd^ buffer. Attention shifts to the yellow letter ‘A’ resulting in its shape and color being stored in the 2^nd^ buffer of DPAAN#1 and DPAAN#2 respectively. Finally attention shifts to the cyan letter ‘T’ resulting it in its shape and color being stored in the 1^st^ buffer of DPAAN#1 and DPAAN#2 respectively. The necessary sequence of buffer and latch activations for such sequential storage is shown in Figure 6A. This is the DPAAN solution to the classic neural binding problem: appropriate gating of attention, relays, and latches ensures that the attributes of a single object are bound together by virtue of them all being represented in similarly indexed DPAAN buffers (Hayworth, 2012).

We propose that something similar happens across the ventral and dorsal cortical visual streams when scenes containing multiple objects are represented and when objects containing multiple parts are represented (Hayworth, 2009). A common attentional ‘spotlight’ in early visual areas ensures that all later areas are representing the attributes of the same object. Individual areas might represent the object’s visual field location, distance, size, orientation, several separate shape characteristics, color, texture, name, and so forth. Each of these highest visual cortical areas would have its own DPAAN that could store the object’s attributes into one of its buffers. As attention shifts sequentially across a set of objects in a scene, a structured representation would be formed across all of these DPAANs, a representation that explicitly binds each object’s shape attributes with its location by virtue of storing these in similarly *indexed* buffers across all the DPAANs. There are many tasks that could benefit from such a structured representation, for example spatial relations processing (Hayworth, 2009), navigational planning, pen and paper arithmetic (Dehaene et al., 1998), and episodic memory.

Figure 7 depicts how such a multi-DPAAN structural representation of a scene can be converted into a single vector representation and stored as a content-addressable declarative memory (DM) using a simple two-level model of the hippocampus. In this model, each DPAAN is provided with its own declarative memory buffer designated ‘DM Level 1’, and then these two ‘DM Level 1’ buffers are themselves tied together via a buffer designated ‘DM Level 2’. Such multi-level funneling is common in some models of associative memory (Sacramento and Wichert, 2011) but in our model we add switchable relays at the inputs and outputs of all levels. As we will show, these allow the DM to be queried in a fill-in-the-blank manner.

**Figure 7.**
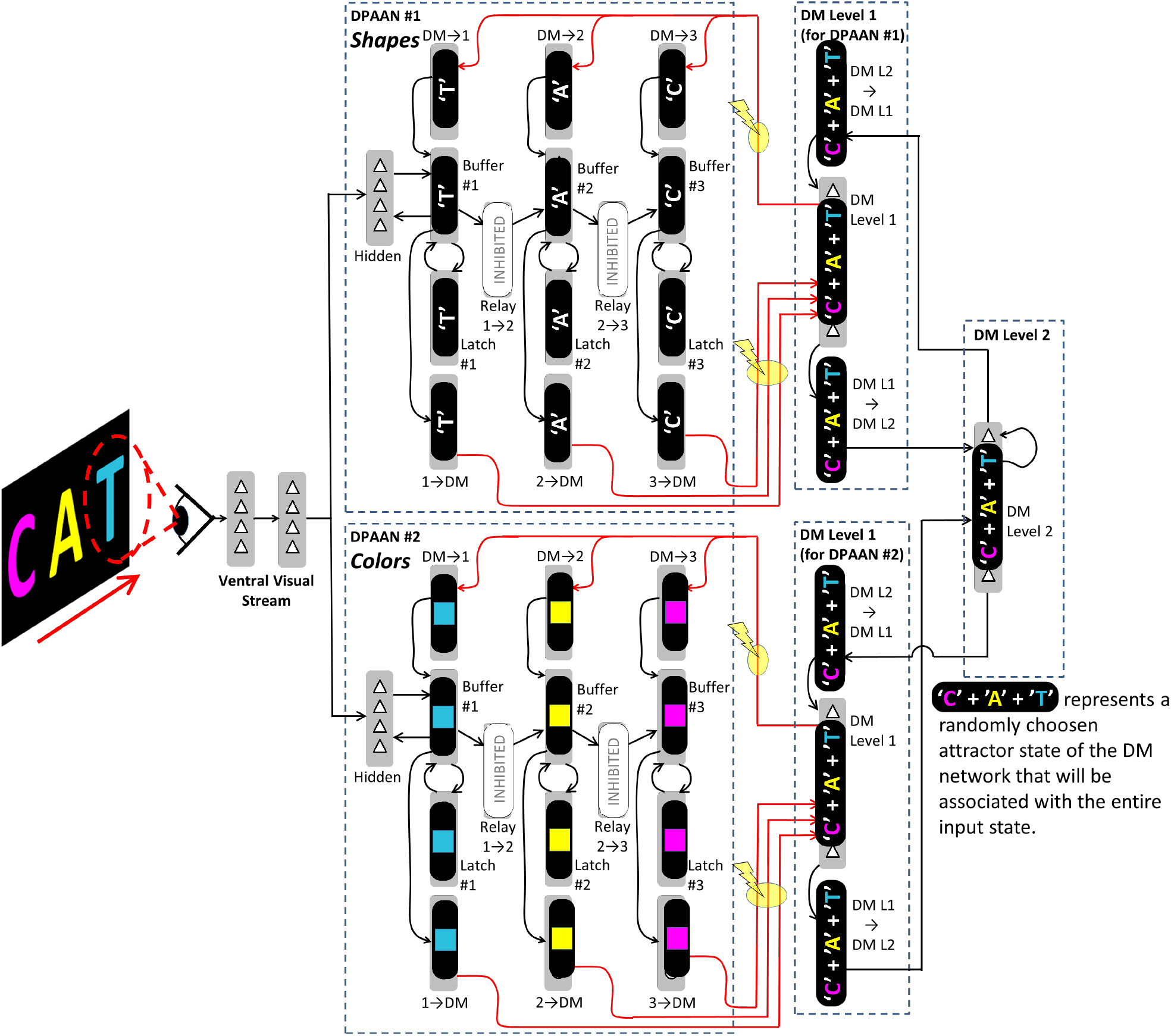
Two DPAAN modules sharing a common input and declarative memory network. Red highlights and lightning bolt symbols show which weight matrices are modified to store a new declarative memory.

#### Setting up DPAAN#1 and DPAAN#2 symbols

As described in previous sections, our model trains the synaptic matrices within each of the DPAANs so that all of its neuronal groups act together as a single autoassociative memory that can subsequently be dynamically partitioned by inhibiting groups of relay neurons. For the current simulation, each DPAAN consisted of three buffers along with a full complement of latches, relays, and equality detection networks, including new relays to and from the DM. We will designate each symbol in DPAAN#1 as 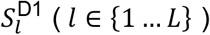 and each symbol in DPAAN#2 as 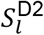. Each of these symbols consists of a randomly generated sparse activation over all of the DPAAN’s neuronal groups. For example, the *l*^th^ symbol in DPAAN#1 consists of a particular activation pattern over all of the following neuronal groups:

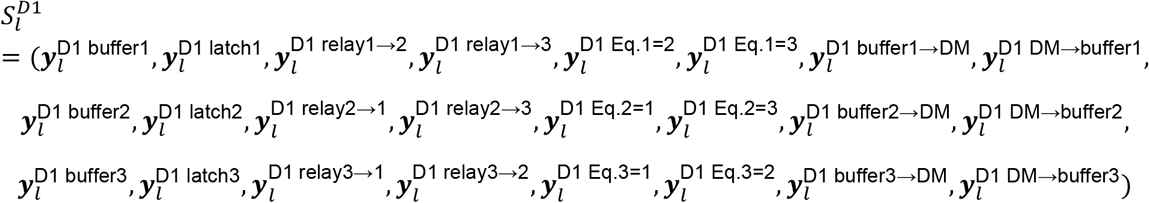

where 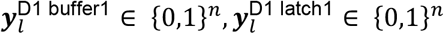, etc., and where there are n = 300 neurons in each group and the sparseness (*α*) of each was set to 0.05 to match the k-WTA dynamics of each group.

DPAAN#2 was set up in a similar fashion:

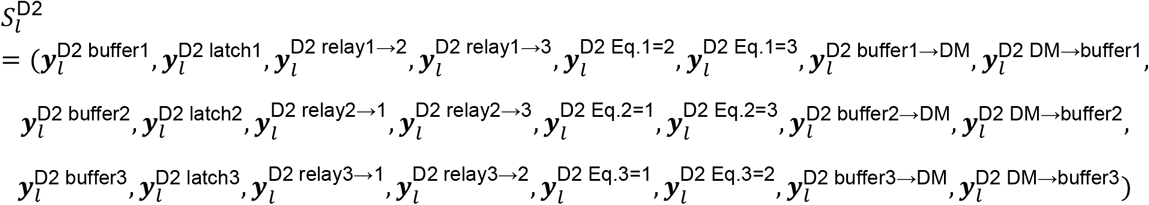

When the DPAANs are representing multiple items as shown in Figure 7, the relays between buffers are inhibited so that buffers 1, 2, and 3 can be locked into different attractor states. Let’s say that DPAAN#1’s symbol *l* = 5 represents the letter ‘C’, symbol *l* = 7 represents the letter ‘A’, and symbol *l* = 9 represents the letter ‘T’. Since DPAAN#2 represents color, let’s say that its symbols *l* = 12, *l* = 14, and *l* = 16 represent ‘magenta’, ‘yellow’, and ‘cyan’ respectively. The activity patterns depicted in Figure 7 would then be:

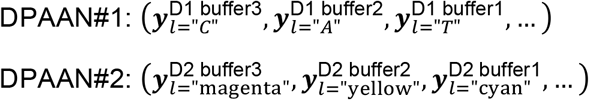

#### Setting up DM symbols

The goal is to have the two level DM network store complex multi-DPAAN patterns such as this by associating each with a single attractor state in the 2-level DM network. In this way an incomplete pattern in the DPAANs can be used to call up the previously associated DM vector which can then be used to reconstitute the rest of the original memory in the DPAANs (For example predicting that the next letter in the sequence ‘C’,’A’, is likely to be ‘T’, or by recalling the colors that were associated with these previously stored letters.)

To accomplish this we first trained the DM network alone so that all of its neuronal groups acted together as a single autoassociative memory. A set of *L* = 100 randomly generated ‘memories’ were created to train the DM network. These memories will be the symbols in the DM that will be associated with patterns in the DPAANs. Each symbol 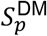 (*p* ∈ {1 … *P*}) consisted of a set of randomly generated sparse activation patterns:

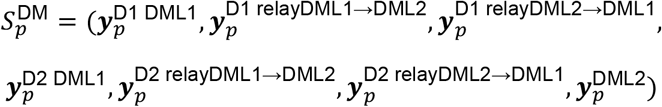

Where 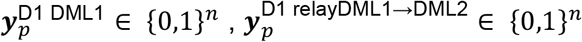, etc. For simulations we used n = 300 and k-WTA sparseness = 0.05. This training of DM symbols was performed the same way the training of DPAAN symbols was, modifying the weight matrices connecting the different neuronal groups in the DM.

#### Storing a declarative memory

At this point each of the DPAANs has been trained with a symbol vocabulary, each of the DPAAN input networks has been trained to classify sensory inputs (e.g. letter shape for DPAAN#1 and color for DPAAN#2), and the DM network has been trained with its own symbol vocabulary, but we have yet to store an actual declarative memory. To actually store a declarative memory a unique pattern of symbols is first latched into the DPAAN buffers, and an ‘unused’ DM symbol, say 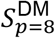, is latched into the DM network. In order to form an association between the DPAAN pattern of symbols and this DM symbol we modify the weight matrices connecting the two. In the simulation the weight matrices to modify are as follows:

Weights from DPAAN#1 to DM: ***W***^D11→DM to D1 DML1^, ***W***^D12→DM to D1 DML1^, ***W***^D13→DM to D1 DML1^
Weights from DM to DPAAN#1: ***W***^D1 DML1 to D1 DM→1^, ***W***^D1 DML1 to D1 DM→2^, ***W***^D1 DML1 to D1 DM→3^
Weights from DPAAN#2 to DM: ***W***^D21→DM to D2 DML1^, ***W***^D22→DM to D2 DML1^, ***W***^D23→DM to D2 DML1^
Weights from DM to DPAAN#2: ***W***^D2 DML1 to D2 DM→1^, ***W***^D2 DML1 to D2 DM→2^, ***W***^D2 DML1 to D2 DM→3^

The lines corresponding to these weight matrices are highlighted in red in Figure 7. These weights are trained like all the others using the cross product rule, but in this case the result is that the entire DPAAN#1 + DPAAN#2 + DM network is trained as a single, multilevel autoassociative memory network.

In summary, whenever it is desired to store the current contents of the DPAANs, an unused DM symbol is loaded into the DM network and the weight matrices highlighted in Figure 7 are trained to store this association.

#### Recalling a declarative memory

Figure 8 shows how specific pieces of a declarative memory can be recalled in a fill-in-the-blank fashion. The letter ‘C’ has been recognized by DPAAN#1’s input network and loaded into its buffer#3, and the letter ‘A’ has been loaded into buffer#2. An attempt to recall what letter comes next in the sequence would involve disinhibiting the relays ***y***^D1 buffer2→DM^ and ***y***^D1 buffer3→DM^ which together drive ***y***^D1 DML1^ into the DM symbol that was associated with the original declarative memory. When the relay 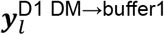 is now also disinhibited this DM symbol drives DPANN#1’s buffer#1 into the correct symbol ‘T’.

**Figure 8.**
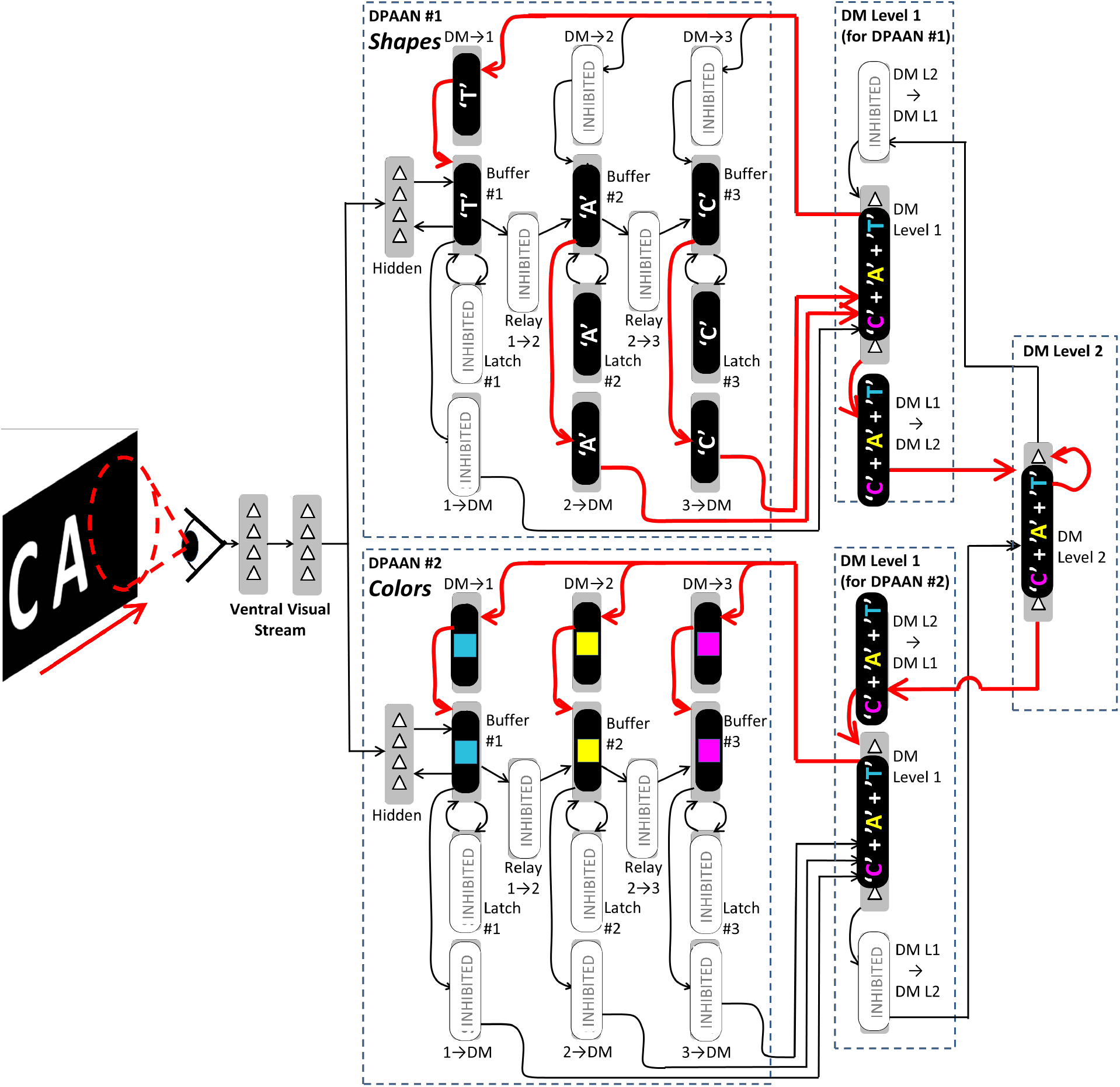
Example of declarative memory recall. This figure depicts the fill-in-the-blank recall of the same declarative memory that was stored previously in Figure 7. Scanning visual attention has loaded DPAAN#1 with the letters ‘C’ and ‘A’. Disinhibiting specific relays (2→DM, 3→DM, DM→1) causes the letter ‘T’ to be recalled and loaded into DPAAN#1’s Buffer#1. Disinhibiting other relays results in the recall of the colors that were previously associated with each of these letters.

Disinhibiting other relays within the DM network and DPAAN#2 allows this recalled declarative memory to fill DPANN#2’s buffers with the stored colors as well as shown in Figure 8.

#### Simulation results

Two DPAANs were trained to have a 100 symbol vocabulary (all DPAAN neuronal groups contained n=300 neurons, 0.05 k-WTA sparseness). A two level DM network was trained to have a 100 symbol vocabulary (all DM neuronal groups contained n=900 neurons, 0.05 k-WTA sparseness). One declarative memory was stored associated with each of these DM symbols for a total of 100 declarative memories. Each memory consisted of assigning a random set of symbols to the six DPAAN buffers. After all memories were stored, recall was tested by latching the correct symbols into two of the DPAAN buffers and simulating network routing and settling (as depicted in Figure 8) to recall the other four symbols and load them into the appropriate DPAAN buffers. Memories storage and recall tests were designed to avoid ambiguous overlapping memories. Using these parameters all DM recall tests were successful.

## Discussion

In this final section, we first summarize our claims and place them in a putative evolutionary context. We then discuss the neuroanatomical and neurophysiological evidence for thalamocortical latch and relay structures of the type our model relies on. Finally, we discuss relationships of our model to ideas in cognitive science, computer science and machine learning.

### Summary and evolutionary perspective

In our model the cortex and thalamus are perhaps best viewed as a collection of DNNs interspersed with recurrent memory buffers that support latching and routing operations. At least in infancy this cortical system is trained and controlled by the BG and other subcortical structures like the SC, but later in life we suggest that this joint system becomes a powerful computational engine seamlessly combining aspects of both symbolic and subsymbolic computation and learning.

A first-order benefit of this joint arrangement would be to supervise-train categorizing DNNs in the cortex, which can in turn provide the BG (via cortico-striatal projections) with more meaningful representations of the external world. A second-order benefit would be the ability to supervise-train deep sensorimotor neural networks within the cortex to better generalize actions that are initially performed by the basal ganglia and brainstem sensorimotor circuits—a model for how cortical consolidation might work (e.g., (Ashby et al., 2010; Pyle and Rosenbaum, 2018)). A third-order benefit could be that the basal ganglia learns to utilize the collection of cortical memory buffers to keep track of the state information important to the current task over an extended period of time, i.e., working memory. A fourth-order benefit could be that the basal ganglia learns to skillfully control storage to, and associative recalls from, longer-term hippocampal and temporal lobe stores. Finally, an exciting fifth-order benefit could be that the system learns to control patterns of routing operations among the collection of cortical buffers so that they can perform complex symbolic computations. Each of these hypothesized benefits incrementally extends the core BG system:

1. **Training of sensory DNNs**—makes BG reinforcement learning more efficient by providing the BG with improved, more generally useful representations
2. **Training of sensorimotor DNNs**—consolidates and generalizes BG sequence learning to allow more complex actions
3. **Control of cortical working memory**—allows the BG’s shallow pattern recognition circuits to learn more complex state-based behaviors
4. **Control of longer-term associative memory storage and recall**—makes it possible for the BG to learn to map its environment and to control complex, temporally-extended behaviors
5. **Control of complex symbolic computations**—allows the BG’s shallow pattern recognition circuits to become syntax-sensitive, allowing rule-based learning and ultimately complex cognition

This list of proposed benefits is meant to suggest an evolutionary progression, where the evolving thalamo-cortical system expands to better serve the needs of a control system initially dominated by the basal ganglia. This hypothetical evolutionary pathway suggests how a DPAAN-like system of gated buffers and relays, as well as contrastive learning procedures, all under control by the basal ganglia, might have emerged incrementally from a more primitive basal ganglia based action control system.

## Neuroanatomical basis of controllable cortico-cortical relays

Several neuro-anatomical structures and pathways are implicated in the selective control of cortico-cortical information flow and in the maintenance of information in working memory.

### Sherman’s “thalamic higher order relays”

Almost all information flowing into the cortex passes through ‘first order’ thalamic relay nuclei. For example, visual information from the retina is transmitted to the primary visual cortex via relay cells in the lateral geniculate nucleus (LGN) of the thalamus. First order relays like the LGN constitute only a small proportion of the volume of the thalamus in primates (Sherman, 2017). In fact, it seems that every cortical region receives projections from and sends projections to the thalamus. This and other evidence led (Sherman and Guillery, 1996) to propose that the majority of the thalamus might constitute a set of ‘higher order’ relays, serving a key role in transmitting information between cortical areas.

For example, the pulvinar nucleus is the main higher order thalamic relay of visual information, projecting to and receiving projections from all areas of the ventral visual stream as well as many other cortical areas (Shipp, 2003). Its relay cell axons project to higher visual cortical areas and terminate in layer 4 in a manner reminiscent of how LGN relay axons drive layer 4 in primary visual cortex (Sherman, 2017). A similar pattern is seen for the projections of other higher order thalamic relays, and tests show these projections drive strong, fast excitatory postsynaptic potentials (EPSPs) in the recipient layer 4 cells (Lee and Sherman, 2008).

What is the driver, in the Sherman and Guillery view, of these proposed higher order thalamic relay cells? Axons from cortical layer 5 pyramidal cells provide a strong ‘driver’ input analogous to the retinal axons driving LGN cells (Sherman and Guillery, 2002). These typically have large synapses, fast ionotropic glutamate receptors, and terminate onto the proximal dendrites of the higher order relay cells (Sherman and Guillery, 2011). In addition, the receptive field properties of these thalamic higher order relay cells appear to mirror those of the layer 5 cortical cells (Sherman and Guillery, 2002).

Crucially for relating these findings with the DPAAN model, there is evidence that these layer 5 projections can originate from a separate cortical area than the one the relay cells are projecting to (Llano and Sherman, 2008; McFarland and Haber, 2002), providing a trans-thalamic route for information to flow between these two separate cortical areas. A study by (Theyel et al., 2010) provided some direct evidence that information critical to cortical computations is being relayed through thalamic higher order relays. They performed optical imaging and whole cell recordings in a mouse thalamocortical somatosensory slice preparation which preserves the primary barrel cortex (S1BF), the secondary somatosensory cortex (S2), and the posterior medial nucleus of the thalamus (POm). The POm is the presumed higher order thalamic relay that connects S1BF to S2. This slice model also preserved the projections of the transthalamic pathway (S1BF → POm → S2) as well as some of the direct cortical projections (S1BF → S2). When the direct axonal projections from S1BF to S2 were completely cut, S2 still showed a robust response to S1BF stimulation. But S2 response was eliminated after the POm relay was ablated.

In summary, there is evidence that higher order thalamic relays provide routes for information transfer among cortical regions, routes operating alongside direct corticocortical projections, and that these routes are necessary for cortical operation (Zhou et al., 2016). Information transferred via these transthalamic relays is modulated by other brain structures, which therefore provide a measure of top-down control over cortical communications. For example, studies have shown that the pulvinar’s connections to higher visual cortical areas play an important role in controlling visual attention (Saalmann and Kastner, 2009), and it is thought that inhibitory projections from the thalamic reticular nucleus (TRN) onto pulvinar relay cells directly modulate information transfer (Lakatos et al., 2016). Specific cortical circuits impinging on the TRN, together with the TRN itself, would then play an ‘operator’ role for this hypothesized thalamic ‘switchboard’ (Kastner et al., 2012; Zikopoulos and Barbas, 2006).

It should be noted that many of the cortico-thalamic projections to higher order thalamic relays are branches of axons that also project to sub-cortical motor centers (Guillery and Sherman, 2011). This has been interpreted to suggest that the messages being transmitted through higher order thalamic relays are “efference copies” of motor output commands, and thus may be involved in sensorimotor integration. Such efference copies have also been suggested to be crucial for reinforcement learning in the striatum (Fee, 2014). Our approach here predicts that thalamic relays would be used more broadly: not only for conveying efference copies of motor commands, but also for routing symbolic information and coordinating supervised training of pattern recognition networks. If implemented via Sherman higher order relays, the information transmitted would be a generalized cortical L5 output, rather than a motor command as such. More generally, the implication that any subcortical projection should be termed a ‘motor output’ copy may merit reconsideration. Many of these projections are to the striatum, for example, which in our model would need the same high-level symbolic information as the cortical routes it is supposed to control.

### The basal ganglia as a potential latch and relay control operator

A subset of higher-order thalamic relays appear to be under direct inhibitory control of the basal ganglia. These basal ganglia-recipient thalamic nuclei are interconnected with the highest levels of sensory cortex (Clower et al., 2005; Middleton and Strick, 1996) and with motor, prefrontal, and “limbic” cortex (Alexander and Crutcher, 1990; McFarland and Haber, 2002). Our model proposes that DPAAN buffers exist in all of these regions. In the highest levels of sensory cortex, DPAAN buffers would store the structured, symbolic representations of sensory inputs and provide a means to supervise-train sensory hierarchies via the clamping of a target at the top-level of the hierarchy.

The output of the basal ganglia contains groups of tonically-active inhibitory neurons which project to brainstem motor circuits and to some thalamic nuclei, normally suppressing their action (Grillner et al., 2005; Grillner and Robertson, 2016). The precise impact on the thalamus of this dis-inhibition of the basal ganglia to thalamus outputs is still the subject of much study, but at least one model suggests that this dis-inhibition could allow cortical driver inputs to pass through the thalamic relay, as in our model (Goldberg et al., 2013).

There are multiple additional potential anatomical candidates for a switchboard that controls the latches and relays. In addition to the basal ganglia acting as a switchboard via topographic disinhibition at its outputs to thalamic relays, other models suggest (Solari and Stoner, 2011) that L1-projecting thalamus constitutes the core switchboard for working memory control. In addition, the claustrum may provide a locus for control of cortico-cortical communication (Solari and Stoner, 2011). In any case, inputs to the thalamus from cerebellum and other structures, aside from the basal ganglia, may also play a role in controlling cortico-cortical information routing.

## Neurophysiological basis of controllable cortico-cortical relays

Our model made a number of neurophysiological assumptions, one of which was the transfer of cortical information through controllable relays. To establish whether such relays may be located in the thalamus, an ideal experiment would simultaneously record from a source and destination cortical regions as well as a thalamic relay linking them. The (Theyel et al., 2010) study described above provided evidence of the thalamus acting as an information-bearing relay. A recent paper (Ito et al., 2015) also argued for the transfer of trajectory-encoding information from the prefrontal cortex through the nucleus reuniens of the thalamus and into the hippocampus, with the nucleus reuniens thus acting as an information-bearing relay.

Another assumption of our model was a multi-part latched memory structure which may comprise a cortico-thalamo-cortical loop. Experimental evidence that persistent firing of prefrontal cortical and associated mediodorsal thalamus neurons underlies working memory can be traced back at least to the studies of (Fuster and Alexander, 1971), and recent evidence continues to support this link (Guo et al., 2017).

A further question is whether such cortico-thalamo-cortical loops are under inhibitory control by the basal ganglia. Evidence for this exists in the songbird, where analogous structures clearly exhibit a basal ganglia controlled loop underlying vocal learning and expression (Goldberg and Fee, 2012). In addition, modeling studies have argued for basal ganglia control of PFC-MD memory loop (Wei and Wang, 2016), and a number of models both utilize a thalamocortical loop as a memory and have the thalamic leg of that loop under inhibitory control by the basal ganglia (Ashby et al., 2005; O’Reilly and Frank, 2006; Vitay and Hamker, 2010).

Another physiological property assumed in our discussion is the use of attractor based coding of symbols, which is crucial to the operation of a DPAAN. Recent evidence in the frontal cortex argues in favor of such a model (Inagaki et al., 2017), as does recent evidence in the hippocampus (Pfeiffer and Foster, 2015), and a variety of important models for, e.g., hippocampal phase precession also rely on this idea (Lisman et al., 2005), although the applicability of the attractor concept in various brain areas is still a matter of debate (Colgin et al., 2010), as are the mechanisms by which attractors may be formed and modified (Koyama and Pujala, 2018; Miconi et al., 2016; Park et al., 2016; Pokorny et al., 2017). Note that our model only proposes attractor-based computation for associative and frontal cortex, not for early sensory areas, which we modeled as trained deep neural networks here.

Finally, our use of Contrastive Hebbian Learning as a synaptic learning rule to train a multi-layer network, in our model of sensory hierarchies, was meant to be illustrative of a wider range of emerging candidate mechanisms for biologically plausible multilayer credit assignment, many of which use some kind of contrastive mechanism for training (e.g., (Guerguiev et al., 2017)), and thus are compatible with our scheme based on clamping of the output buffer with target patterns. Some of these mechanisms are reviewed in (Marblestone et al., 2016).

## Alternative models of thalamic information routing

The thalamus is a heterogeneous structure and our understanding of its anatomy and of its roles in working memory and control of cortical information flow is still preliminary (Granger, 2007; Halassa and Kastner, 2017). Indeed, there exist multiple conceptually distinct models of the functioning of cortico-thalamo-cortical relays as it relates to anatomy and physiology.

Some models of higher order thalamic relay nuclei differ from Sherman’s view, by suggesting that the flow of represented information is purely cortico-cortical, but that the thalamus modulates donor and/or recipient cortical areas to selectively control this cortico-cortical information flow. For example, (Ketz et al., 2015) suggest that “thalamic circuits may play a functional role in selectively gating communication between cortical regions by *synchronizing* them in various frequency bands … information is transmitted via the cortico–cortical connections to the next cortical region or regions, while the [higher order] thalamic nuclei *selectively activate the appropriate downstream cortical area* that will be engaged in the next level of processing”. Similarly, (Schmitt et al., 2017) argues for “thalamic control of functional cortical connectivity …. which is dissociable from categorical information relay”. This viewpoint is particularly well developed in connection with thalamic areas controlling attention, e.g., the pulvinar (Saalmann et al., 2012; Womelsdorf and Fries, 2007) (though there are also a variety of non-synchrony-based models of attention (Bobier, 2011)), where it is suggested that the thalamus provides modulatory drive to local inhibitory interneuron circuits in the cortex to generate increased synchrony and other attentional effects such as normalization control (Buschman and Kastner, 2015). Some theory has explored how such synchrony-based modulation of cortico-cortical communication might be integrated into a basal ganglia controlled architecture (Pouzzner, 2017). In such a scenario, our model would need to be modified, and additional aspects would need to be explained; for instance, whether thalamus-induced synchrony between cortical areas could enable transfer of attractor states as required in DPAAN.

In another model of thalamo-cortical working memory and its control structures (Solari and Stoner, 2011), the role of Sherman higher order relays is de-emphasized. Instead, each region of cortex is reciprocally connected to a corresponding area of “specific thalamus” via a L6 to thalamus to L3 loop, which maintains activity in persistent thalamo-cortical attractor states, in a manner conceptually similar to our model of working memory (the thalamic reticular nucleus plays a supporting role here by helping to ensure excitatory/inhibitory balance). Instead of Sherman’s L5 to thalamus to L4 relays, this paper emphasizes an information-bearing role for L6 projections to thalamus, which Sherman considers to be modulatory in nature (Sherman, 2017). The authors of (Solari and Stoner, 2011) interpreted their model not in terms of categorical / representational information relay but in terms of control flow in a “confabulation” model of attractor state control (somewhat analogous to our latching mechanism but without a direct analog of our relays), but the anatomy itself might potentially support both interpretations. Furthermore, in (Solari and Stoner, 2011), some regions of thalamus send projections to L1 cortex, which are taken as control inputs for the recipient cortical areas, with a function somewhat related to our proposed “latching” mechanisms for working memory. Yet other thalamic domains, such as intralaminar thalamus, send thalamo-cortical projections that are taken to selectively gate the outputs of L5 pyramidal neurons, serving as a mechanism for control of behavioral output (Solari and Stoner, 2011). Other papers are suggestive of a “driver” function for thalamo-cortical projections to L1 (Garcia-Munoz and Arbuthnott, 2015), including in basal ganglia recipient thalamic areas. Thus, several anatomical structures in the corticothalamic system, other than those emphasized by the models of Sherman and Guillery, might potentially support controllable memory buffers, latches and relays of the type we have proposed^3^.

## Relationships with cognitive science concepts

If a DPAAN-like system of linked, controllably latched and routed memory buffers indeed exists in the brain, it would have several key implications for our understanding of cognitive architecture. In particular, we propose that DPAAN allows the implementation of a “brain wide production system” reminiscent of discrete production system models of cognition like ACT-R (Anderson et al., 2004), SOAR (Laird and Newell, 1996) and others. As overviewed in Figure 9, the basal ganglia could learn, via reinforcement learning, a set of IF-THEN rules that allow it to control, via the thalamus, the routing of information into and out of memory structures, thereby implementing a production system. Crucially, these IF-THEN rules could be syntax sensitive (by being sensitive to DPAAN equality detection networks) and could manipulate syntax (by instigating buffer-to-buffer coping operations and fill-in-the-blank declarative memory recalls). The DPAAN architecture was, in part, motivated by the goal to show how these types of syntax sensitive IF-THEN rules, which are at the heart of cognitive architectures like ACT-R, could be learned by a shallow pattern recognition network like the one assumed to exist in the striatum (Hayworth, 2012).

**Figure 9.**
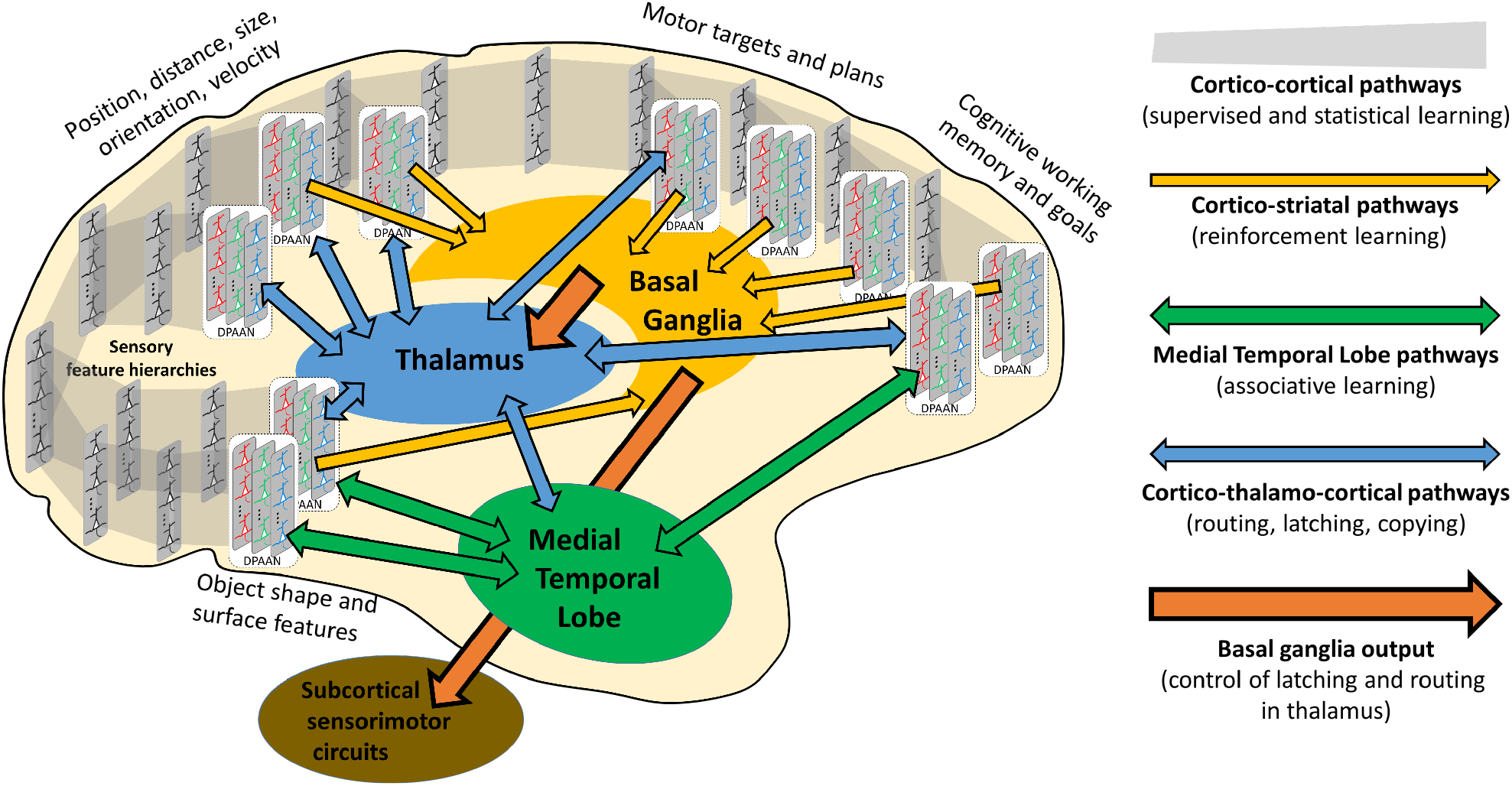
System-level sketch of how cortical buffers and thalamic latches and relays might interact to form a brain-wide production system. The most fundamental behaviors of the animal are handled by **subcortical sensorimotor circuits** whose instigation and sequencing are controlled via inhibitory projections from the **basal ganglia**. **Cortico-striatal projections** provide meaningful state information to the basal ganglia to guide action choices (Redgrave et al., 1999). The reinforcement-trained pattern recognition circuits in the basal ganglia can be thought of as implementing a set of if-then ‘production rules’. The basal ganglia, via its projections to the **thalamus**, can also learn to control latching and routing in the cortex itself by disinhibiting specific **cortico-thalamo-cortical pathways**. A key goal of such latching and routing is to direct the storage of task-relevant information in **medial temporal lobe** (MTL) structures like the hippocampus, allowing this state information to be recalled to both direct future behavior and, central to our proposal here, to provide supervised training of **cortico-cortical projections**. This is accomplished by latching recalled states into specific cortical buffers thereby driving some form of Contrastive Hebbian Learning (CHL) in the cortex. We suggest that CHL via the clamping of cortical buffers underlies the training of cortical sensory hierarchies, the training of cortical sensorimotor streams, and the consolidation (i.e. transfer) of declarative and procedural memories from the sites of their initial learning (MTL and Basal Ganglia respectively) to cortex. Evolution of the flexible control of latching and routing in the cortex would also allow for collections of cortical buffers to be trained as Dynamically Partitionable AutoAssociative Networks (DPAANs). Because these DPAANs provide for buffer-to-buffer copying and equality detection operations, they fully support the types of symbolic/syntactic processing assumed by classic production system architectures like ACT-R for example (Anderson, 2007), but do so in a way that allows syntax-sensitive if-then production rules to be implemented by shallow reinforcement-trained pattern recognition circuits in the basal ganglia (Hayworth, 2012). (Diagram inspired by (Cisek and Kalaska, 2010)).

As discussed in the introduction, our proposed model of information transfer between latched buffers, containing corresponding pieces of shared attractor-state based symbols, also provides an implementation of the much-discussed concept of “variable binding”, reminiscent of early proposals for anatomical variable binding (Hadley, 2009; Marcus, 2001) and related to the more recent proposal of (Legenstein et al., 2016) which is based on linked neural assemblies. Many of the most successful cognitive science models implemented within the ACT-R framework rely heavily on such variable binding (Anderson, 2007), and our DPAAN-based model offers one potential way that these models might be anatomically grounded.

In addition, these same control structures could mediate aspects of supervised learning in the sensory hierarchies, which we modeled as deep neural networks trained via contrastive supervision and credit assignment mechanisms. We have illustrated the use of latched buffers and relays to orchestrate the supervised deep training of a feedforward hierarchy, taken to crudely model visual input streams.

Yet this same system could also drive the training of learned connections for other purposes. For example, trainable cortical-cortical associative connections could link the DPAAN buffers directly, bypassing the proposed trans-thalamic relay route. Under appropriate conditions, these cortico-cortical associative connections could be trained to predict or mimic the “symbolic” operations taking place in the DPAAN buffer system. Thus, “symbolic” buffer-to-buffer transfer, recall, and comparison operations within the cortico-hippocampal system, under step by step control by the basal ganglia, could become “consolidated” into associative linkages inside the cortex. Whole series of such “symbolic” operations could also be “chunked” into a single step in this manner, a process referred to as ‘production compilation’ in ACT-R (Anderson, 2007). Thus, the model would have potential implications for consolidation (Kumaran et al., 2016) of both procedural and declarative memories.

## Relationships with computer science concepts

This paper has hypothesized that certain aspects of the brain’s anatomy and physiology can be conceptualized as implementing controllable routing of information among a set of cortical memory buffers. At least on the surface, our proposed model is somewhat analogous to the controlled routing of information that underlies modern digital computers. In this subsection, we explicitly explore this analogy to see how far it can be taken and, importantly, to point out areas where the analogy breaks down. Figure 10A depicts a digital computer’s routing circuitry, and Figure 10B depicts the analogous situation in our proposed model of cortico-thalamo-cortical routing in the brain. The juxtaposition of these two diagrams should help make clear the analogous elements that we describe here.

**Figure 10.**
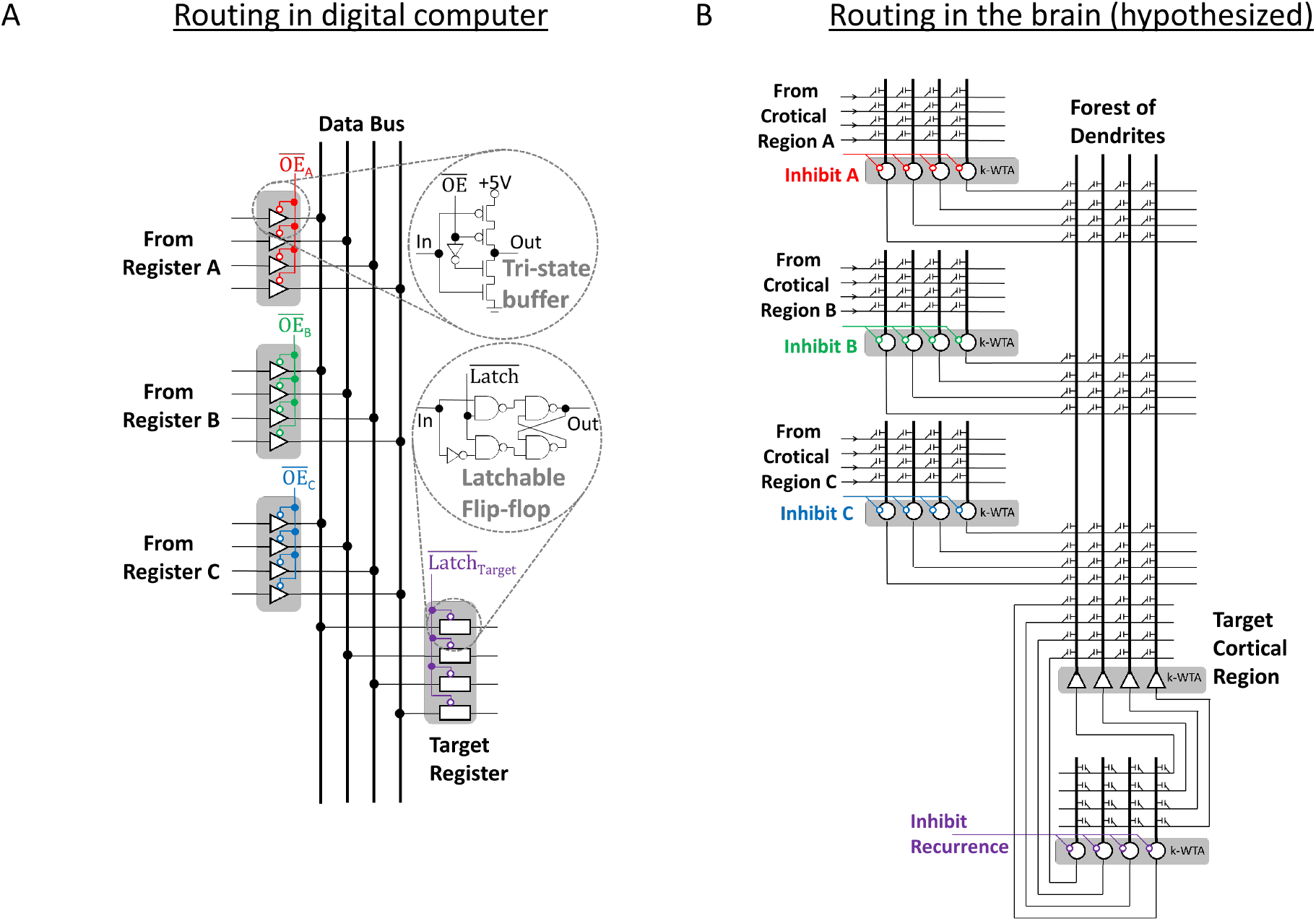
Analogy to computer architecture concepts. (A) Electric circuit diagram showing how tristate buffer circuits and flip-flops are used to route information along a digital computer’s data bus. (B) Analogous diagram showing how routing of information is performed in our model.

A digital computer routes information between a source register and a target register by means of a data bus—a collection of wires to which all registers are connected. The computer’s control system ensures that only one register is allowed to drive the data bus at any one time. To accomplish this, the outputs of all registers are connected to the bus via tri-state buffers—a particular arrangement of transistors which can be electronically commanded into a high-resistance state. A bus routing operation is started by setting the 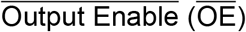 line of the source register’s tri-state buffers to ‘Low’, electronically signaling the tri-state buffers to put that source register’s voltages on the data bus. Analogy: the computer control system’s enabling of a collection of tri-state buffers is analogous to the basal ganglia disinhibiting a thalamic higher order relay.

Digital computers store individual bits of information (a ‘1’ or a ‘0’) by means of a circuit called a ‘flip-flop’. A flip-flop contains two cross-connected NAND gates which together form a bistable circuit—if the top NAND gate’s output is ‘High’ then the bottom NAND gate’s output is forced to be low and vice versa. Additional circuit elements provide a means to momentarily disrupt this bistable circuit when the flip-flop’s 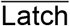 line is brought ‘High’, forcing the bistable circuit into a state determined by the flip-flop’s input line. When the 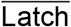 line is returned to ‘Low’ this new state is ‘latched’ in and will remain for as long as power is supplied to the circuit. Analogy: The trained excitatory connections and pooled inhibitory connections (implementing k-WTA) within our hypothesized cortico-thalamo-cortical bidirectional associative memory buffer make it behave like a ‘multidimensional flip-flop’, i.e. a physical system having many stable attractor states. Transiently inhibiting the thalamic half of a cortico-thalamo-cortical memory buffer allows it to be ‘latched’ into a new state reflecting its current inputs.

In order to route (copy) digital data from a source register to a target register in a digital computer, the source register’s 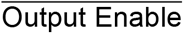 line is first brought ‘Low’ putting its voltages on the data bus. Then the target register’s 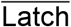 line is momentarily brought ‘High’ (allowing the data bus’s voltages to force the target register’s flip-flops into matching states), and then ‘Low’ again (latching these new bit states in place). Such register-to-register copy operations require precise synchronization of source and target registers, synchronization that is provided in digital computers by global control and clock signals. Analogy: We hypothesize that the thalamocortical oscillations observed in mammalian brains (Ketz et al., 2015) reflect analogously synchronized transfers of information among cortical regions. This puts the observed synchronies in a different perspective: rather than synchrony causing improved communication among areas (Ketz et al., 2015; Pouzzner, 2017; Saalmann, 2014), the observed synchrony is a result of underlying latch and relay operations.

There is a very important place where this computer analogy breaks down: The flip-flops within each digital register behave independently of all other flip-flops allowing each N-bit register to be latched into any arbitrary binary state out of the 2^N^ possible. Furthermore, digital computers are designed with precise wiring enabling the one-to-one transfer of bits between registers and one-to-one comparison operations. That is, the nth bit of every register is wired to the same (nth) wire of the data bus. In contrast, neuronal connections in a biological brain are assumed to be only crudely determined by genetics—regional targets are specified but local synaptic connections are initially random prior to learning. Thus a biological brain cannot rely on built-in one-to-one wiring either for memory storage or for information routing.

It is seldom appreciated, but it is precisely this one-to-one wiring that allows a digital computer to act as a physical symbol system (Fodor and Pylyshyn, 1988) complete with variable binding operations. Each variable in a computer program is associated with a particular memory register and each symbol in a program’s vocabulary is associated with a particular pattern of ones and zeros.

The brain, in contrast, lacks such one-to-one wiring, but the DPAAN mechanism solves this problem by introducing an attractor-based anatomical variable binding mechanism (Hayworth, 2012). By training all of a DPAAN’s buffers, relays, and equality detection networks together, a common set of global attractor states is formed. The symbol vocabulary of each part of this network simply consists of pieces of these global attractor states. This training ensures that there is a one-to-one relationship between buffers, not at the individual neuron level but at the individual *attractor state* level. And it is this learned one-to-one relationship between the attractor states of separate buffers that allows copy and comparison operations in the DPAAN and that ultimately allows for variable binding and syntax sensitive rule applications.

## Relationships with machine learning

### DPAAN and supervised learning

We have made a specific claim about where at least some of the “learning signals” or “error signals”, for at least some forms of deep learning (i.e., multilayer credit assignment) might arise in the brain. We propose that the final layer of each specific cortical hierarchy (e.g., face area, object area, visual word form area, etc.) has an associated region of thalamus, such that the two form a bidirectional associative memory (BAM) buffer. We propose that there are innate circuits in the thalamus and midbrain that will trigger this BAM to latch under appropriate situations. When latched to a particular “symbol”, the buffer contents can be used in a contrastive supervised learning process to train the upstream sensory hierarchy to produce that symbol from a broader set of inputs.

This allows the creation of a ‘cost function’ built from examples of the form: ‘this percept should be giving rise to this symbol activation’ (in the declarative case, while a similar idea can be used for sensorimotor procedural consolidation). Contrastive learning will minimize this cost function as long as the synaptic weights between the latched BAM and the earlier parts of the sensory hierarchy can be modulated (perhaps through oscillations) to give rise to a (-) phase and a (+) phase that each synapse can respond to.

For example, once a symbol for the face of “mom” is established, the system could latch the face buffer under rotations of the head in order to generalize the range of inputs that give rise to that symbol and train the visual hierarchies accordingly. Moreover, the hippocampus can reinstate the BAM with a previous representation based on cross-modal information (i.e., your mother says ‘cup’ for a new object and this allows you to clamp in the same object representation for ‘cup’ that you associated with a previously viewed but visually dissimilar exemplar).

This BAM can also be triggered to latch by disinhibition of the basal ganglia’s outputs to the thalamus. Because the basal ganglia’s control policy is learned via dopamine-mediated reinforcement learning, this brings the ability to decide *when* to do supervised training, and with *which* clamped target patterns, under reinforcement learning control. In other words, the animal can reinforcement learn when and how it is appropriate to supervise-train a specific cortical area. This would allow the brain to have flexible, learnable control over when to group percepts into the same category, and for this control to be driven by reinforcement-learned policies in the basal ganglia or by innate heuristics present in other sub-cortical structures.

Thus, we have defined a biologically implementable cost function (vectorial distance to BAM-latched symbol) for supervised learning, how it is minimized by gradient descent (contrastive learning implemented by modulating the weights between the BAM and the rest of the hierarchy, as well as within the rest of the hierarchy), who does the training (basal ganglia controlled thalamic latches and relays) and how the training is bootstrapped from limited data (by innate subcortical circuits, by reinforcement learning in the basal ganglia, and by hippocampal recall). (Of course, the cortical circuitry may also be trained by other, unsupervised cost functions, e.g., based on prediction (O’Reilly et al., 2017), or by reinforcement (Chubykin et al., 2013).)

This framework predicts that the dimensionality of control that the supervised training system used by the brain can exert is limited to the number of independent basal ganglia output channels and other independent sub-cortical inputs to thalamic relays and latches. It suggests that distinct cortical regions (Vul et al., 2012) (e.g., a face area vs. a word form area) could be defined by being subjected to distinct training signals or cost functions, which would in turn be defined by connectivity with distinct thalamic latches and relays, which would be addressed differently by basal ganglia output structures and other sub-cortical inputs. The particular pattern of latch and relay operations to which each buffer in a higher-level cortical area is subjected, by the basal ganglia and other regions, would determine a region-specific training procedure for it and its immediately upstream regions. This is distinct from other suggested frameworks for how such distinct cortical areas would arise, e.g., based on distinct transformation invariance properties (Leibo et al., 2011).

We speculate that such an architecture might have been ‘discovered’ by evolution first as a means to bootstrap the learning of more meaningful cortical representations feeding the striatum, but eventually led to a system that could support symbolic computation, i.e., variable binding and production-like rules. In particular, the latching and relay mechanisms might have originally developed via selection for improvements in mechanisms for training stimulus generalization in the sensory hierarchies, but then adapted towards discrete control of memory buffers and more generalized routing.

### Future directions for the integration of the DPAAN framework with machine learning

Memory-augmented neural networks (Graves et al., 2016) and relation networks (Santoro et al., 2017) have recently been demonstrated as a means of integrating symbolic operations with trained artificial neural networks. In both those cases, the system is fully differentiable and end-to-end backpropagation based training is used to define the weights in the network (although recent works have started to depart from end-to-end training of full architectures (Wayne et al., 2018), the memory structures themselves often remain differentiable). In our DPAAN model, in contrast, attractor-based computation and gated control of information flow in attractor networks are used to perform symbolic operations, as well as to train a sub-network via a Contrastive Hebbian approximation of backpropagation. In turn, the policy for gating of buffer and relay states is proposed to be learned by reinforcement learning in the basal ganglia. Our system also stores and recalls information from an episodic memory crudely based on the hippocampus. Of course, the brain may not be an end-to-end differentiable system, and thus our model can be viewed as a proposal for a non-differentiable “outer loop” of infrastructure (itself trained in part via reinforcement learning in the basal ganglia) in which training of sub-networks can occur. At least one recent paper within artificial intelligence has espoused a similar idea (Garnelo et al., 2016), though without specifying a fully neural implementation.

Our proposal for basal ganglia control of information routing in prefrontal cortical networks may also have implications for models of “learning to learn” in the joint prefrontal cortex basal ganglia system, e.g., for models in which plasticity mediated reinforcement learning in the basal ganglia gives rise to an activity-mediated learning system in the prefrontal cortex (Wang et al., 2016, 2018). In our model, the PFC would not only serve as an input to basal ganglia action selection systems, but would also be subject to latch and relay control by the basal ganglia outputs; this might introduce additional possibilities for how an activity-mediated learning system in the PFC could be trained.

Recent advances in deep learning have also shown how constraining higher layers (later stages) of processing to support symbolic operations (e.g., relational reasoning) can influence, through backpropagation, the earlier stages of processing, forcing them to produce “object-like” outputs that serve as good substrates for the symbolic manipulation (Santoro et al., 2017). We speculate that structuring or constraining deep learning models to have a DPAAN-based output layer and outer control loop could similarly drive the formation of learned upstream representations that are useful in the context of these downstream routing and memory structures, e.g., driving the learned representations to be useful for DPAAN-based symbolic processing.

While DPAAN based models have not yet begun to be developed for high performance on machine learning tasks, we suggest that cross fertilization of these ideas with existing approaches in structured machine learning architectures could be an intriguing future direction to inform both fields.

## Acknowledgements

We thank Konrad Kording, Greg Wayne, Gary Marcus, Chris Eliasmith, Wolfgang Maass, Soren Solari, Dario Amodei and Bragi Lovetrue for helpful discussions, along with the participants of the Kavli Futures Symposium: Towards a taxonomy of cortical computations.

## Footnotes

3 It should also be noted that our proposed implementation of cortico-thalamic bi-directional associative memories, containing strong “driver” connections in both directions, would conflict with Crick and Koch’s “no strong loops” hypothesis (Crick and Koch, 1998) for cortical and cortico-thalamic connectivity. However, this hypothesis was motivated based on the concept that such loops could lead to uncontrolled oscillations, whereas in our model, these loops are under tight inhibitory control by local circuits, the basal ganglia, and other inputs, which would serve the same purpose.

